# Growth, metabolism, and the benefits of clock-enabled anticipation

**DOI:** 10.64898/2026.10.01.755848

**Authors:** Michele Monti, David K. Lubensky, Pieter Rein ten Wolde

## Abstract

Circadian clocks are found in myriad species across the tree of life. They are commonly assumed to provide a selective advantage by enabling organisms to anticipate daily environmental changes. It has proven remarkably difficult, however, to move beyond this general idea to explain precisely what advantage anticipation confers and to quantify how large this benefit is. Here, we address these challenges by developing a mathematical model of diurnally varying growth and metabolism in the cyanobacterium *Synechococcus elongatus*, which fixes carbon through photosynthesis during the day and stores a portion of it as glycogen for use during the night. Specifically, we extend the framework introduced by Hwa and coworkers (Scott *et al*., Science **330**, 1099 (2010)), which quantifies the relationship between growth and proteome composition, to a circadian context. We find that the fastest growth is achieved when glycogen production during the day is minimized to the amount required to sustain essential nighttime processes; growth then occurs exclusively during the day, in agreement with experimental observations. However, the slow relaxation of the proteome, arising from the intrinsically slow growth rate of *S. elongatus*, severely constrains the implementation of this optimal strategy: In the absence of anticipation, the cell must grow at night in order to produce proteins essential for nighttime metabolism. In contrast, with anticipation enabled by a circadian clock, the bacterium can instead adjust its proteome late in the day, allowing it to avoid less efficient nighttime growth altogether, thereby significantly enhancing its growth rate.

---

Organisms ranging from cyanobacteria to plants face the challenge that they can acquire crucial nutrients only at certain times of day. For example, obligate photoautotrophs cannot fix inorganic carbon in the absence of sunlight. As a result, these lifeforms need to store nutrients during one part of the 24h diurnal cycle for use during other parts. Notably, most such organisms possess a circadian clock, which allows them to anticipate daily changes in their environment and to adjust their metabolism ahead of time. Yet, it remains unknown what the best strategy is for adjusting metabolism to periodic, diurnal environmental changes or how much faster organisms employing this strategy can grow than those without a clock. Here, we address these questions through a computational model of the cyanobacterium *Synechococcus elongatus*.

*S. elongatus* is among the most studied organisms that exhibit circadian rhythms. Its primary source of carbon is CO_2_, which it fixes during the day via photosynthesis. Intriguingly, while the bacterium predominantly grows during the day, it also stores a fraction of its carbon in the form of glycogen, which is then consumed during the night [1, 2]. In light-dark (LD) cycles, a diurnal rhythm of glycogen accumulation is observed for both wild-type cells and clock null mutants, while under constant light (LL) conditions this rhythm is observed only in wild-type cells [1, 2]. The glycogen rhythms are thus under the control of the circadian clock but can also be directly induced by the environment.

These physiological rhythms are mirrored by rhythms in gene expression and protein levels driven by the circadian clock [3– The expression of the majority of rhythmic genes peaks near either subjective dawn or subjective dusk [4, 5]. In particular, the microarray data of [5] revealed that genes underlying the Calvin cycle and glycogen synthesis predominantly peak around dawn while genes of the Oxidative Pentose Phosphate Pathway (OPPP) and those involved in breaking down glycogen mainly peak at dusk [2].

Why cyanobacteria store glycogen remains a subject of debate. In LD, cells do not grow appreciably at night, indicating that glycogen is not used primarily to fuel growth. One possibility is that stored glycogen allows cells to protect themselves against Reactive Oxygen Species (ROS) [2, 8], which are a by-product of photosynthesis [9]. Consistent with this idea, a mutant that is deficient in circadian gene regulation rapidly accumulates ROS during the day that cannot be cleared at night [8]. Moreover, the main reductant for ROS detoxification is NADPH rather than NADH [9]; the OPPP, which is the main channel for glycogen consumption at night, produces NADPH, while glycolysis generates NADH [2, 8]. In addition to this use of NADPH to scavenge ROS, the OPPP is also needed at the end of the night to prime rapid initiation of photosynthesis at daybreak [10], and its resources can be required early in the night to complete DNA replication [11].

While these and related experiments [12, 13], together with large-scale computational modeling studies [14–16] provide detailed information about how physiology and gene expression change over the 24 h period, many questions remain open: First and foremost, can we quantify the benefit of the clock? Classic experiments clearly show that there is an advantage to having a clock whose period matches the environment but do not directly measure how large this benefit is or explain how the clock confers it [17, 18]. In general, anticipation becomes beneficial when the cellular response is slow [19, 20]. In the absence of active protein degradation, the timescale for the relaxation of the proteome is given by the growth rate [21–23], which in *S. elongatus* tends to be low, with doubling times typically of order 10 hours or longer [24–26]. This suggests that anticipation should be valuable to the cyanobacterium. But how much does anticipation increase the growth rate, and what trade-offs determine the optimal timescale for anticipation?

To address these questions, we employ the framework developed by Hwa and coworkers for quantifying the relationship between growth and proteome composition [22, 23, 27–31]. In this framework, the system is not described via detailed signaling and metabolic networks, but rather via coarse-grained protein sectors, which facilitates a quantitative mathematical analysis. While previous studies have successfully applied this framework to the bacterium *Escherichia coli*, we here extend it to study circadian metabolism of cyanobacteria.

Importantly, we focus here on the benefits of anticipation enabled by an internal timekeeper that is assumed to be accurate and noise-free. We do not address the fascinating but separate question of whether a true, autonomous clock or a so-called hourglass is a more effective timekeeper [32–35]. As we are interested primarily in *S. elongatus*, which does have a true clock, we often refer to our timekeeper as a clock.

Specifically, we consider a sequence of three closely related models: the *quasi-equilibrium model*, the *slow-proteome model*, and the *anticipation model*. These models share the same minimal features that capture the interplay between using carbon to drive growth during the day and storing carbon as glycogen during the day to fuel processes at night; these include a ribosome sector that synthesizes proteins to allow growth, a carbon sector that controls the rate of carbon fixation, and a carbon-storing sector that shunts some of the fixed carbon into glycogen during the day. All three models also have an OPPP sector that consumes glycogen to generate NADPH [2, 8] and meet the cell’s physiological needs at night. The models differ in how quickly protein levels can adjust to environmental changes and in whether the cell uses a clock to anticipate such changes. By comparing the models’ behavior, we are thus able to quantify how much faster cells can grow when a clock allows them to anticipate and to understand the mechanisms behind this growth rate increase.

We find that, in the presence of the clock, the cells adopt an extreme strategy, in which they exclusively grow during the day. Two factors underlie this strategy: first, producing enzymes that do not directly contribute to growth, i.e. the glycogen-storing enzymes, lowers the growth rate; and, second, setting aside carbons as glycogen for nighttime growth means that they only contribute to growth with a delay, and this is less efficient than using them immediately to catalyze more carbon fixation and protein synthesis. Cells should thus store only as much glycogen as they need to survive during the night, without saving any extra for growth. However, this strategy can only be implemented when a clock is present to allow the cell to upregulate the OPPP sector before nightfall. Without such anticipation, in the absence of active protein degradation the cell must either make the OPPP proteins, and thereby grow, at night or express them throughout the day, even when they are not needed. The central benefit of the clock is thus that it allows the proteome to adjust late in the day to prepare for the night, eliminating the need for less efficient nighttime growth.

A prediction of our theoretical analysis is that clocks are only valuable when the cell needs significantly different proteome compositions in daytime and nighttime. To demonstrate that this is indeed true of *S. elongatus*, we conclude by comparing our model’s results to proteomics data [7]. To this end, we present a new method, Principal Mass Harmonics Analysis (PMHA), which allows us to quantify how much protein mass is redistributed between day and night. It is based on projecting trajectories in protein composition space onto the ellipses specified by their lowest time harmonics. Applying this analysis to the data of [7] shows that about 8% of the protein mass is redistributed between day and night. Our model can reproduce this shift, but also predicts that at higher growth rates this can increase up to nearly 25%.

## I. THEORY

Throughout this paper, we assume that cells are grown in an environment with alternating 12h periods of light and dark (LD 12:12), so that the day has a total length *T* = 24h, and that the light intensity is constant throughout the 12h light periods. We take it to be light (i.e. daytime) for 0 *< t <* 12h = *T/*2, where *t* is the time, and dark (nighttime) for 12h = *T/*2 *< t <* 24h = *T*.

### A. Protein sectors

The central ingredients of our model are coarse-grained protein sectors and the balance of fluxes between them (Fig. 1) [22, 23, 27–31]. A sector can be thought of as a set of coordinately regulated proteins that together perform some basic function like carbon fixation. Each protein belongs to one and only one sector, so that the total protein mass *M* in the cell is the sum of the masses *M*_*α*_ in the different sectors *α*: *M* = Σ_*α*_ *M*_*α*_. Correspondingly, we may define the mass fraction *ϕ*_*α*_ of a sector as *ϕ*_*α*_ = *M*_*α*_*/M*.

**FIG. 1:**
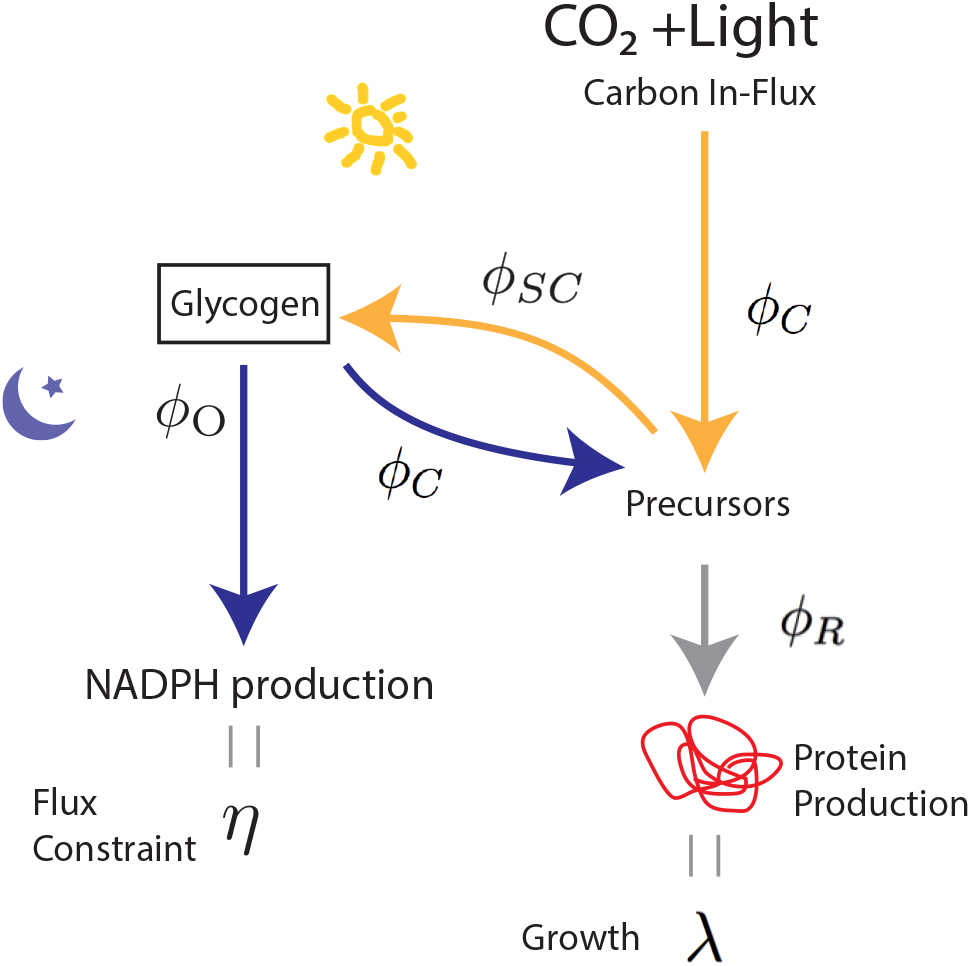
Overview of our model. During the day, the source of carbon is CO_2_, which is fixed via photosynthesis. The carbons are processed via the carbon sector *ϕ*_C_. A fraction of carbon is stored in the form of glycogen via the carbon-storing sector *ϕ*_SC_; the other part is delivered to the ribosomal sector *ϕ*_R_ to drive growth with growth rate *λ*. During the night, a fraction of glycogen is funneled into the OPPP sector to generate NADPH with flux constraint *η*; the remaining fraction is channeled via the carbon sector *ϕ*_C_ into the ribosomal sector *ϕ*_R_, to potentially drive growth.

We are here mainly interested in the flux of carbon atoms. A minimal model therefore contains a ribosome sector with mass fraction *ϕ*_R_ = *M*_R_*/M*, responsible for protein synthesis, and a carbon sector with mass fraction *ϕ*_C_ = *M*_C_*/M* that fixes carbon and incorporates it into amino acids. These sectors must ultimately be defined empirically [31]. In particular, the C-sector consists of those proteins that are upregulated in response to a carbon limitation, yet downregulated in response to translation inhibition. The carbon limitation could take the form of reduced CO_2_ and light levels during the day or of reduced glycogen levels during the night. We anticipate that this sector contains not only the photosynthesic machinery, but also enzymes of the TCA cycle and those responsible for degrading glycogen. Similarly, the R sector contains those proteins upregulated in response to translation inhibition but downregulated in response to carbon limitation and is expected to consist primarily of ribosomal proteins.

In *S. elongatus*, the main channel for glycogen consumption during the night is the OPP pathway [2, 8]. Our model therefore includes an OPPP sector with mass fraction *ϕ*_O_. The OPPP runs parallel to glycolysis, and it may seem natural to assume that this sector contributes to generating building blocks for amino-acids, as the carbon sector *ϕ*_C_ does. However, while the OPPP is active at night, the cells do not grow at night, suggesting that this sector is not coupled to the ribosomal sector. Its main function, it appears, may be to generate NADPH, a reductant that scavenges ROS [2, 8].

The next sector contains enzymes that store carbon as glycogen during the day; its fraction is denoted by *ϕ*_SC_. Enzymes belonging to *ϕ*_SC_ include glycogenin and glycogen synthase, which are upregulated during the day in *S. elongatus* and other cyanobacteria [2, 6, 36, 37]. Expressing the glycogen-storing enzymes is expected to slow down growth during the day (since glycogen storage does not contribute to daytime growth) but potentially enables growth at night.

In keeping with previous work on *E. coli* [27, 28, 31], our model finally includes a fixed unresponsive fraction *ϕ*_Q_. Together, the C, R, Q, SC, and O sectors represent the minimal ingredients needed to capture circadian metabolism in a photosynthetic organism. It is of course possible that these sectors could split into smaller sectors in some circumstances—for example, the C-sector might be divided into separate sectors for photosynthesis and glycogen breakdown. Ultimately, more experiments will be needed to determine whether any such additional subdivisions are important for describing the phenomenology of circadian growth.

### B. Quasi-equilibrium model

We next present a simple limiting case of our full model. In general, the sector fractions *ϕ*_*α*_ will respond to changes in nutrient availability arising from shifts between day and night and from the gradual consumption of stored glycogen during the night. In the absence of active protein degradation, this response happens on a timescale that is set by the growth rate *λ* [21– In *S. elongatus*, however, this timescale is typically not much shorter than the ~12h scale over which nutrient influxes vary [25, 38–40]. Thus, the proteome composition will in general lag behind changes in nutrient availability. Below we will take this slow relaxation of the proteome into account. However, to introduce the main elements of the model, it will be instructive first instead to assume that the growth rate is so high that the proteome is always in quasi-equilbrium with respect to the instantaneous nutrient levels set by the CO_2_ and light levels during the day and by the glycogen level during the night. In this limit, the growth rate during the day is simply a constant; at night, it depends implicitly on time through time-varying glycogen levels. We call this model the quasi-equilibrium model.

Experiments by Hwa and coworkers on *E. coli* show that the steady-state growth rate varies linearly with the size of the R and C sectors [27, 28, 31]. These linear relationships can be understood by combining the following ideas, which we also adopt in our quasi-equilibrium model: a) in steady-state the carbon fluxes *J*_*α*_ through the different sectors *α* = R, C are balanced, so that there is no build-up of intermediates like amino-acids; b) the growth rate *λ* is proportional to the flux through the sectors; and c) the flux through a sector scales linearly with the size of the sector, as would be expected, for example, if the sector consists of enzymes that are operating close to saturation. Our model incorporates the three ingredients a)-c), but adds two additional elements: d) during the day, a certain fraction of the flux through the carbon sector is shunted into glycogen synthesis; and e) during the night, some glycogen is funneled into the carbon sector and some into the OPPP. Importantly, only the flux through the carbon sector flows into the ribosome sector and hence contributes to growth; the nighttime flux through the OPPP is important for generating NADPH but cannot, in our model, drive growth.

#### 1. Flux balance and growth rate

The first two ingredients a) and b) from the preceding paragraph imply that in the quasi-equilibrium model the growth rate *λ*(*t*) is given by

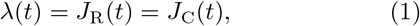

where we have absorbed proportionality factors into the the definitions of the fluxes *J*_R_ and *J*_C_. Observation c) means that the flux through the ribosomal sector takes the form

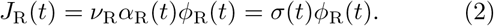

Here, the coefficient *ν*_R_ describes the translational efficiency of an active ribosome, and the quantity *σ*(*t*) = *ν*_R_*α*_R_(*t*) is the overall, instantaneous translational efficiency, taking into account that only a fraction *α*_R_(*t*) of the ribosomes is active at any given instant (as discussed in more detail below) [23].

The ingredients c) – e) similarly imply that the flux through the C sector that flows into the R sector is given by

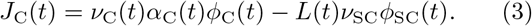

Here, *L*(*t*) is an indicator function that is 1 during the day and 0 during the night. The first term on the right-hand side describes the flux through the carbon sector. During the night, all of this flux flows into the ribosome sector to potentially drive growth. During the day, however, a certain fraction of the flux through the carbon sector is not used for growth but rather devoted to storing glycogen; this is described by the second term. We note that the nighttime flux through the OPPP sector does not enter this expression, because in our model the OPPP never contributes directly to growth by fuelling the ribosome sector. The factor *α*_C_(*t*) in the first term of Eq. 3 is the fraction of the carbon sector that is active, and the parameter *ν*_SC_ in the second term describes the efficiency of the glycogen-storing enzymes and is taken to be constant. Similarly *ν*_C_(*t*) in the first term is a measure of the efficiency of the carbon sector. It depends on the quality of available carbon sources and on how efficiently a given mass of C sector protein can convert these sources into protein building blocks [28, 31]. For the photosynthetic organisms of interest here, during the day the principal carbon source is CO_2_, which means that the value of *ν*_C_ will depend strongly on light levels and also on the concentration of CO_2_. Since we model the light intensity as a step function, during the day 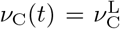 is constant, where the superscript L indicates a quantity that applies when it is light, i.e. daytime. In contrast, during the night, the principal source of carbon is stored glycogen, which is used up as the night progresses. This affects the carbon-processing efficiency. We will model this as

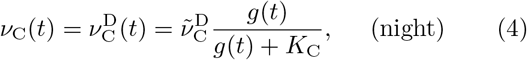

where *g*(*t*) is the ratio of total glycogen mass to total protein mass in the cell and is proportional to the time-dependent concentration of glycogen, *K*_C_ is the glycogen level at which the enzyme efficiency is halved, and the superscript D indicates a quantity assocated with the dark, i.e nighttime. The parameter 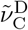 is the maximal efficiency of the carbon sector with glycogen as the carbon source; it does not depend on time.

#### 2. Inactivating growth

Experiments have shown that *S. elongatus* does not grow at night [1, 2]. We thus hypothesized that the cyanobacterium must be able to inactivate some or all of its ribosome and carbon sectors at night, to prevent them from consuming glycogen to drive growth when that is not desired; this is consistent with recent experimental findings [41]. The parameters *α*_R_(*t*) and *α*_C_(*t*) (Eqs. 2 and 3) allow us to systematically study the effect of inactivating these sectors. To limit the number of parameters, we make the simplifying assumptions for the quasi-equilibrium model that i) the fractions are the same, *α*_R_(*t*) = *α*_C_(*t*) = *α*(*t*), and ii) that this fraction switches between *α*(*t*) = *α*^L^ = 1 during the day and a constant value *α*(*t*) = *α*^D^ during the night. Experiments indicate that after a nutrient upshift the translational efficiency *σ*(*t*) of *E. coli* rapidly approaches the new steady-state value set by the new environment [22, 23, 42], suggesting that the assumption of an instantaneous change upon a day-night shift is reasonable [21, 23].

Combining all five ingredients a) - e) with these simplifications, i.e. Eqs. 1-4, yields (wth *β* = L, D)

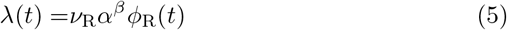

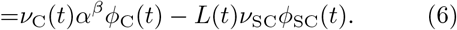

#### 3. Proteome balance

It follows from the fact that Σ_*α*_ *M*_*α*_(*t*) = *M* (*t*) that the protein mass fractions always satisfy

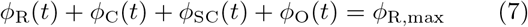

where *ϕ*_R,max_ ≡1 − *ϕ*_Q_ is the maximal ribosomal fraction, which is achieved when *ϕ*_C_ = *ϕ*_SC_ = *ϕ*_O_ = 0.

#### 4. OPPP flux constraint

Following [2, 8], we assume that the cyanobacterium must produce a certain amount of NADPH via the OPPP to meet its minimal nighttime physiological needs, e.g. to fully scavenge the ROS produced during photosynthesis. To ensure that this occurs, we demand that the integrated flux through the OPPP during the night take a fixed value *η*:

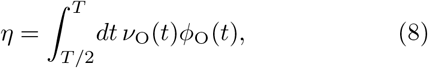

where *ν*_O_(*t*) describes the activity of the OPPP enzymes, which in our model are only active during the night. This activity depends on the glycogen level:

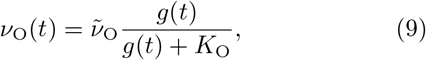

where 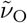 is the maximum activity and *K*_O_ the glycogen concentration at which the activity is halved. This integral constraint is the simplest form that ensures that some glycogen is stored during the day and consumed at night, as observed [1, 2]; without it, the fastest growth would be achieved by setting *ϕ*_SC_ = *ϕ*_O_ = 0 and completely shutting down all metabolic processes at night (see Modeling Results below).

#### 5. Growth rate in quasi-equilibrium model

In the quasi-equilibrium model, the highest growth rates are attained when *ϕ*_SC_ is zero during the night and *ϕ*_O_ is zero during the day. For simplicity, we further assume that 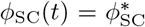 is constant during the day and that 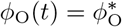 is constant during the night. Given values for these two parameters and for *α*^D^, it is straightforward to solve Eqs. 5-7 for *ϕ*_R_, *ϕ*_C_, and *λ*, yielding the instantaneous growth rate for the quasi-equilibrium model:

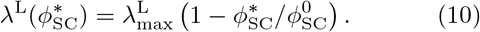

Here, 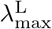 is the maximum growth rate during the day,

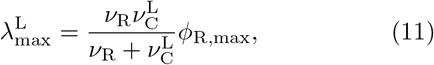

which is reached when no glycogen is stored and *ϕ*_SC_ = 0. The quantity 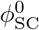 is the size of the carbon-storing sector at which the growth rate goes to zero. It is given by

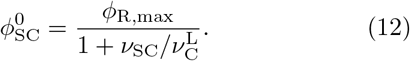

Similarly, we find for the growth rate during the night

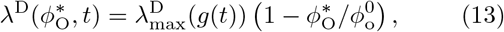

with

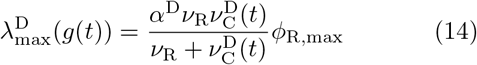

and 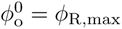.

A few points are worthy of note. First, Eqs. 10 and 13 show that the growth rate decreases linearly with *ϕ*_SC_ and *ϕ*_O_, respectively. Scott *et al*. derived a similar relation for the growth rate when an unnecessary protein is expressed [27]. This highlights the idea that storing glycogen and driving the OPPP reduces the growth rate because these processes require proteins, *ϕ*_SC_ and *ϕ*_O_, that do not directly contribute to growth. Second, building a carbon reservoir during the day is necessary to meet the OPPP flux constraint during the night (Eq. 8). And while diverting carbon flux to this reservoir reduces the instantaneous growth rate during the day (Eq. 3), the stored glycogen can fuel growth at night (Eqs. 13 and 14). Third, we see that the maximum growth rate during the night depends on the fraction of ribosomes that are then active, *α*^D^.

#### 6. Glycogen reservoir dynamics

Glycogen is stored during the day at a rate *ν*_SC_*ϕ*_SC_ (Eq. 3) and consumed at night for growth, at a rate proportional to 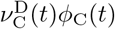 (Eqs. 3 and 4), and for the OPPP pathway, at a rate proportional to *ν*_O_(*t*)*ϕ*_O_(*t*) (Eqs. 8 and 9). The glycogen level thus obeys

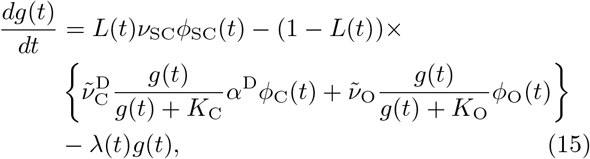

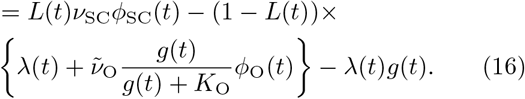

The term *λ*(*t*)*g*(*t*) describes the effect of dilution because of the cells’ exponential growth at rate *λ*(*t*). In the last line we have exploited the fact that in the quasiequilibrium model the growth rate at night equals the flux through the carbon sector; clearly, faster nighttime growth consumes more glycogen.

### C. Slow-proteome model

The quasi-equilibrium model assumes that the proteome is always in equilibirum with respect to the instantaneous nutrient levels, but this will not in general be the case. In this section, we thus remove the assumption that the protein sector mass fractions *ϕ*_*α*_(*t*) relax infinitely fast and develop a model that instead includes their full time evolution, with a characteristic timescale set by the grow rate *λ*.

The basic outline of the flux-balance formalism with time-dependent growth rates is well-established [23]. We first write the mass fractions of the different protein sectors as

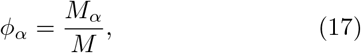

where *M*_*α*_ is the mass of sector *α* = R, C, SC, O, Q and *M* is the total protein mass of the cell. The total rate at which proteins are synthesized is given by

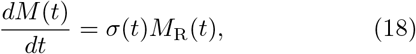

where *M*_R_ is the mass of the ribosomal sector and *σ*(*t*) = *ν*_R_*α*_R_(*t*) is the overall, instantaneous translational efficiency introduced below Eq. 2. When we divide the above equation by *M* (*t*), we obtain the instantaneous growth rate [21, 23]

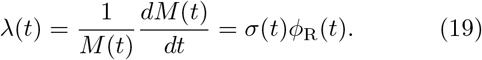

To understand the evolution of the different sectors, we denote the fraction of ribosomes that are allocated to making protein sector *α* as *χ*_*α*_. The sector mass *M*_*α*_ then obeys

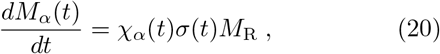

and the proteome fraction obeys [21, 23]

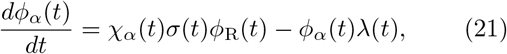

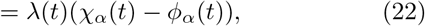

where in going to the last line we have exploited Eq. 19. In Eqs. 20-22, we assume that protein concentrations relax primarily via dilution, as commonly observed in bacteria [43–45]. We can see from Eq. 22 that, because of this assumption, the growth rate *λ* sets the characteristic timescale for changes in the mass fractions *ϕ*_*α*_.

Eqs. 21 and 22 still leave open the questions of how the growth rate *λ*(*t*) and the allocation fractions *χ*_*α*_(*t*) are set. Both of these questions are more subtle than in the quasi-equilibrium model: Whereas in quasi-equilibrium the fractions *ϕ*_*α*_ are chosen so that the fluxes through the C and R sectors balance, in the slow proteome model the time evolution of the *ϕ*_*α*_ must instead follow Eq. 22, and flux balance will not generally be achieved without additional regulation. We must thus choose which sector sets the overall growth rate *λ*(*t*) and how to regulate the other sector to prevent large fluctuations in levels of small molecule intermediates. Similarly, it is implicit in the quasi-equilibrium model that ribosomes are allocated such that new proteins are synthesized in proportion to the existing mass fractions *ϕ*_*α*_, so that these fractions are maintained at steady state. But outside of steady state, there is no comparable constraint on the values of the *χ*_*α*_’s. Here, we propose a set of simple rules to address these issues and rationalize these rules on purely heuristic, phenomenological grounds; in the *Supporting Information*, we illustrate a plausible mechanistic basis for the same rules by showing that they may be derived as a limiting case of the qORAC model previously introduced in the context of metabolic control in non-photosynthetic microbes like *E. coli* [46].

To motivate our rules for setting *λ*(*t*) and the *χ*_*α*_(*t*), we note that the cell needs to meet several challenges: First, upon a change in the environment—like a transition from from day to night—that changes the coefficients *ν*_C_ and *ν*_R_, it must quickly adjust the activities *α*_R_ and *α*_C_ to balance the fluxes through the C and R sectors; failure to do so would lead to a rapid depletion or buildup of intermediates like free amino acids (see *Supporting Information*). Second, on longer time scales, it should adapt the protein mass factions *ϕ*_*α*_ so that, in a constant environment, they eventually approach the steady state fractions from the quasi-equilibrium model. And third, ideally it should meet these challenges while maintaining the highest possible growth rate. How might this be accomplished?

A control strategy that meets all three objectives is one that always maximizes the strength of the weakest link, i.e. that maximizes the flux through the sector that limits the growth rate. To explain this strategy, we note that the C and R sectors each can sustain a maximal flux, which is reached when the proteins in that sector are fully active. To meet the above objectives, we therefore operate the sector with the smaller maximal flux at full capacity, while maintaining flux balance by downregulating the activity, and thus the *α* value, of the sector with the larger maximal flux. The model thereby maximizes the instantaneous growth rate in changing conditions.

Concretely, we thus take the growth rate to be the smaller of the two maximum fluxes through the two sectors,

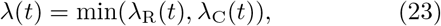

where the maximum allowed fluxes are

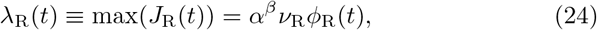

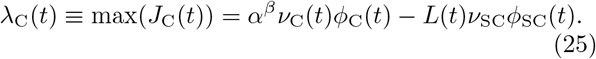

In this scheme, the limiting sector is maximally active, with an activity *α*^*L*^ = 1 during the day and an activity equal to a parameter *α*^D^ to be optimised during the night. To maintain flux balance, the sector that is not limiting then must be further down-regulated through a time-dependent factor *α*_R,C_(*t*) *< α*^*β*^ chosen so that the fluxes through the two sectors are equal.

Having given a prescription for determining the growth rate, we now turn our attention to the ribosome allocation fractions *χ*_*α*_(*t*). As in the quasi-equilibrium model, we assume that the SC and O sectors have target levels 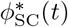 and 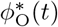, where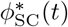 has a constant non-zero value 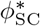 during the day and is zero at night, and 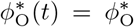 is non-zero at night, but zero during the day. We set the ribosome allocation fractions equal to these target levels, 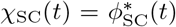 and 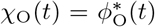, so that *ϕ*_SC_(*t*) and *ϕ*_O_(*t*) would relax to their target levels at long times in a constant environment; because of the slow relaxation of the proteome, however, they will not in general actually reach these values during a 12h light or dark period. We also set *χ*_Q_(*t*) = *ϕ*_Q_ so that *ϕ*_Q_ remains at the same constant value.

With these values of *χ*_*α*_ given, the fraction of ribosomes remaining for the R and C sectors is

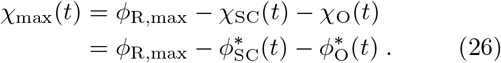

Following our arguments above, we propose to allocate these ribosomes via a strategy akin to a bang-bang controller that seeks to reach the maximal, steady-state growth rate in a fixed environment as quickly as possible. We hence give all of the available ribosomes to the limiting sector. That is,

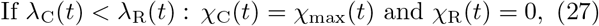

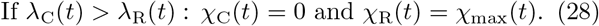

In a fixed environment this allocation rule reaches the steady state *λ*_R_ = *λ*_C_ in a finite time, in contrast to alternative strategies that exhibit exponential relaxation to steady state, which is only truly attained in the infinite time limit.

Finally, in Eq. 25, *ν*_C_(*t*) during the night, i.e.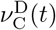, depends on the glycogen level (see Eq. 4), which varies with time according to the same dynamics as in the quasiequilibrium model (Eq. 16). We also continue to demand that the net flux through the OPPP pathway at night equal a prescribed value *η*. The dynamics of the slow-proteome model is thus fully specified by Eqs. 22-28, together with Eqs. 8, 9 and 16.

### D. Anticipation model

In both the quasi-equilibrium model and the slow-proteome model, all changes in parameter values and model behavior occur at the moment when day switches to night or night switches to day. In other words, neither model makes any use of the ability afforded by a circadian clock to *anticipate* upcoming nightfall or daybreak.

Because the proteome relaxes with a timescale set by the growth rate *λ* (Eq. 22), which is typically comparable to or slower than the 24h circadian period, such anticipation might be expected to allow significantly faster growth. To understand how much faster, in this section we introduce the anticipation model, which generalizes the slow-proteome model by allowing parameter changes to occur an anticipation time *T*_a_ before the actual changeover between day and night.

Specifically, with the first subjective day running from *t* = 0 to *t* = 12h, in the anticipation model *χ*_O_(*t*) is equal to the nonzero constant 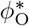 from *t* = 12h *− T*_*a*_ + (24h)*k* to *t* = (24h)(*k* + 1) *− T*_*a*_, where *k* = 0, 1, 2, …, and is zero otherwise. Similarly, *χ*_SC_(*t*) is equal to 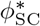 from *t* = (24h)*k − T*_*a*_ to *t* = 12h + (24h)*k − T*_*a*_ and is zero for other times. The governing equations are otherwise the same as in the slow proteome model, which corresponds to the special case *T*_*a*_ = 0. Other rhythmic parameter changes—like the switches between 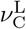and 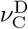and between *α*^L^ and *α*^D^—continue to occur exactly at nightfall and daybreak without any anticipation. Similarly, although the enzymes of the OPPP sector can now be produced ahead of time, they become active and begin consuming glycogen only during the night. This might suggest that these parameters are directly light-dependent, but for the purposes of our model they might equally well be clock-regulated with a different phase from *χ*_O_(*t*) and *χ*_SC_(*t*). To limit the optimization space, we assume that the O and SC sectors have the same anticipation time *T*_*a*_ and that both O and SC sector proteins are actively produced for 12h of each 24h cycle; we will see that this is already sufficient for anticipation to confer a real benefit. It will also turn out that there is negligible nighttime growth for optimal parameters, so that the values of the *χ*_*α*_’s at night have little effect. The main effect of anticipation is thus to shut down production of SC sector proteins and to initiate production of O sector proteins late in the day.

### E. Parameter Values

Except where we state that we optimized over or varied a particular parameter (e.g., see below, *α*^D^ or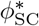), all of our simulations were performed at a single set of parameter values chosen to be biologically reasonable and consistent with key available data on *S. elongatus*. In particular, we chose 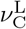 to reflect high light levels capable of generating a doubling time of around 6 hours in constant light (LL) conditions [25] and set parameters related to the SC and O sectors so that the average dry mass fraction of glycogen is roughly 10% [47]. We also set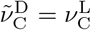so that a given flux of carbon has the same ability to generate growth at night as during the day. A full description of how we estimated parameter values and a table of these values is included in the *Supporting Information*.

## II. MODELING RESULTS

We would like to know the maximum attainable growth rate for each of our three models, from which we will be able to determine how much anticipation can increase growth. But what decides which rates are attainable? Whereas parameters like the *ν*’s are essentially set by the environment or by biochemical kinetics, we imagine that the cell can readily adjust the three parameters *α*^D^, 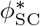, and 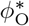 (e.g. Eqs. 10-14 for the quasi-equilibrium model) through regulatory processes. The cell must still reach a given integrated OPPP flux *η* (Eq. 8), so these three parameters cannot be varied independently. Moreover, we show in the *Supporting Information* that the growth rate depends more weakly on 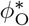 than on the other two parameters, so we usually take 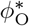 to be fixed. For each model—and for each value of *T*_*a*_ for the anticipation model—we thus maximize the average growth rate over 24h, ⟨*λ*⟩, as a function of *α*^D^ and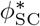, subject to the constraint of Eq. 8.

### A. Quasi-equilibrium model

We begin by examining the quasi-equilibrium model, which allows us to identify a central design principle. Fig. 2A shows a heat map of the time-averaged growth rate ⟨*λ*⟩ as a function of the fraction of proteins that store glycogen during the day, 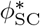, and the nighttime active fraction *α*^D^ of the C and R sectors (see Eqs. 5 and 6). The heatmap also includes contour lines of the net flux *η* through the OPPP sector (Eq. 8). The gradient of *η* is in the direction of increasing 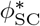 and decreasing *α*^D^: increasing the former allows the cells to store more glycogen during the day, which tends to increase the flux through the OPPP at night; decreasing the latter slows down growth at night, which means that less glycogen is needed to fuel growth and more is funneled into the OPPP sector, again increasing *η*. Importantly, along a given contour line of *η*, the growth rate is maximized for *α*^D^ *→* 0 (Fig. 2A, inset): the optimal strategy is to completely inactivate the C and R sectors at night, so that there can be no growth, and thus only to grow during the day.

**FIG. 2:**
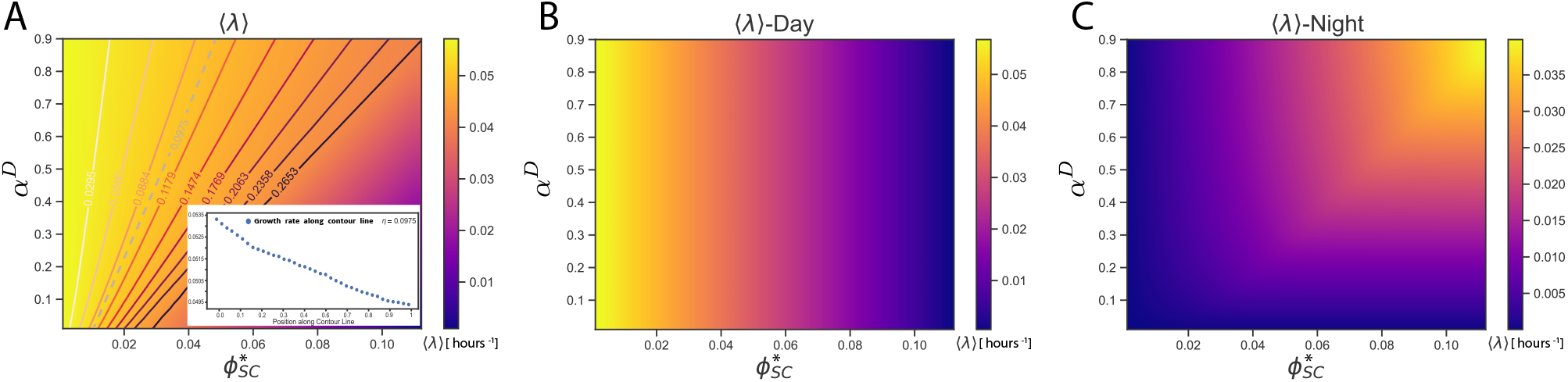
Optimal growth strategy in the quasi-equilibrium model. In this model, the proteome fractions relax instantaneously to quasi-equilibrium with the instantaneous levels of glycogen, *g*(*t*). The instantaneous growth rate is given by Eqs. 10 and 13, and the reservoir dynamics by Eq. 16. (A) Heat map of the average growth rate over day and night, 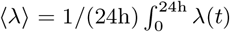, as a function of the active fraction *α*^D^ of the C and R sectors at night (see Eqs. 5 and 6) and the target size 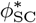 of the carbon-storing sector (which, in the quasi-equilibrium model, is equal to the actual mass fraction *ϕ*_SC_). Projected onto the heatmap are contourlines of constant flux *η* through the OPPP at night (Eq. 8). (B) Heat map of the average growth rate during the day. (C) Heat map of the average growth rate during the night. Along a given contour line of constant *η* the average growth rate ⟨*λ*⟩ over 24 hrs is maximized for *α*^D^ → 0, as illustrated by the inset in panel A, which gives the growth rate along the contour of constant *η* = 0.0975, where the coordinate along the contour (horizontal axis) runs from 0 to 1 from lower left to upper right: the optimal strategy that maximizes the growth rate over 24 hrs is to grow exclusively during the day. Parameter values in *Supporting Information*.

Why is it beneficial to grow exclusively during the day? As panels B and C of Fig. 2 show, increasing the target fraction 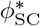 of glycogen-storing enzymes lowers the growth rate during the day but also allows the cell to store more glycogen, which enables it to grow faster at night. However, the loss in daytime growth always exceeds the gain in nighttime growth. One reason is that SC sector enzymes do not enhance the carbon flux devoted to growth. Yet, because the *ϕ*_SC_ sector is small compared to the other sectors, this effect is fairly small.

There is also a second effect: putting aside carbon that is fixed during the day for growth at night lowers the average growth rate. Essentially, by delaying converting carbon into protein, the cell wastes some of this carbon’s capacity to catalyze more growth. To illustrate this effect, it is instructive to compare two extreme cases, one in which the cell only grows during the day, and one in which it only grows at night. For simplicity, let *ν*_R_, *ν*_O_, *ν*_SC_*→∞*, so that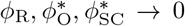. The cost of expressing *ϕ*_SC_ described above is then absent, and the total protein mass is given by the mass *M*_C_ of the carbon sector. During daytime, this mass then grows as *ν*_C_*M*_C_ = *dM*_C_*/dt*, and, with initial mass *M*_C,0_ at daybreak, the mass at the end of the day is 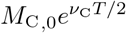; since the cell does not grow at night in this scenario, this is also the mass after *T* = 24 h. In contrast, if the cell only grows at night, then during the day all carbon is devoted to making glycogen, and the total glycogen mass *M*_G_ obeys *ν*_C_*M*_C_ = *dM*_g_*/dt*; since the cell does not grow during the day, *M*_C_ remains constant, and at the end of the day the mass of accumulated glycogen is *M*_G_(*T/*2) = *ν*_C_*M*_C,0_*T/*2. During the night this is all converted into protein mass, so after 24 hrs the mass is *M*_C,0_ + *ν*_C_*M*_C,0_*T/*2. This is always lower than 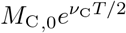 Growing during the day makes it possible to capitalize on the power of exponential growth: any increase in mass is immediately used to further increase the uptake of additional mass.

The central prediction of the quasi-equilibrium model is thus that cells prefer not to grow at night, as observed experimentally [48]. However, the quasi-equilibrium model assumes that the proteome relaxes instantly, while the actual relaxation rate, in the absence of protein degradation, is set by the growth rate, and so is fairly low for cyanobacteria [25, 26]. If the cell’s proteome cannot adapt quickly to day-night shifts, can it still manage to grow primarily during the day, while also having a large enough OPPP sector to meet its needs at night? To address this question, we next turn to the slow proteome model.

### B. Slow-proteome model

Fig. 3A shows the heat map of the average growth rate as a function of the activity fraction *α*^D^ and the glycogen-storing sector target fraction 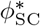 for the slow-proteome model. A comparison to the heat map of the quasi-equilibriun model (Fig. 2A) makes clear that the slow relaxation of the protein sectors dramatically reduces the growth rate. The contour lines of constant OPPP flux *η* have also changed qualitatively. Along each contour line there is now an optimal, non-zero, value of *α*^D^ that maximizes the average growth rate over 24 hrs ⟨*λ*⟩. Clearly, in the slow-proteome model it is necessary to grow at night, in stark contrast to the behavior of the quasi-equilibrium model.

**FIG. 3:**
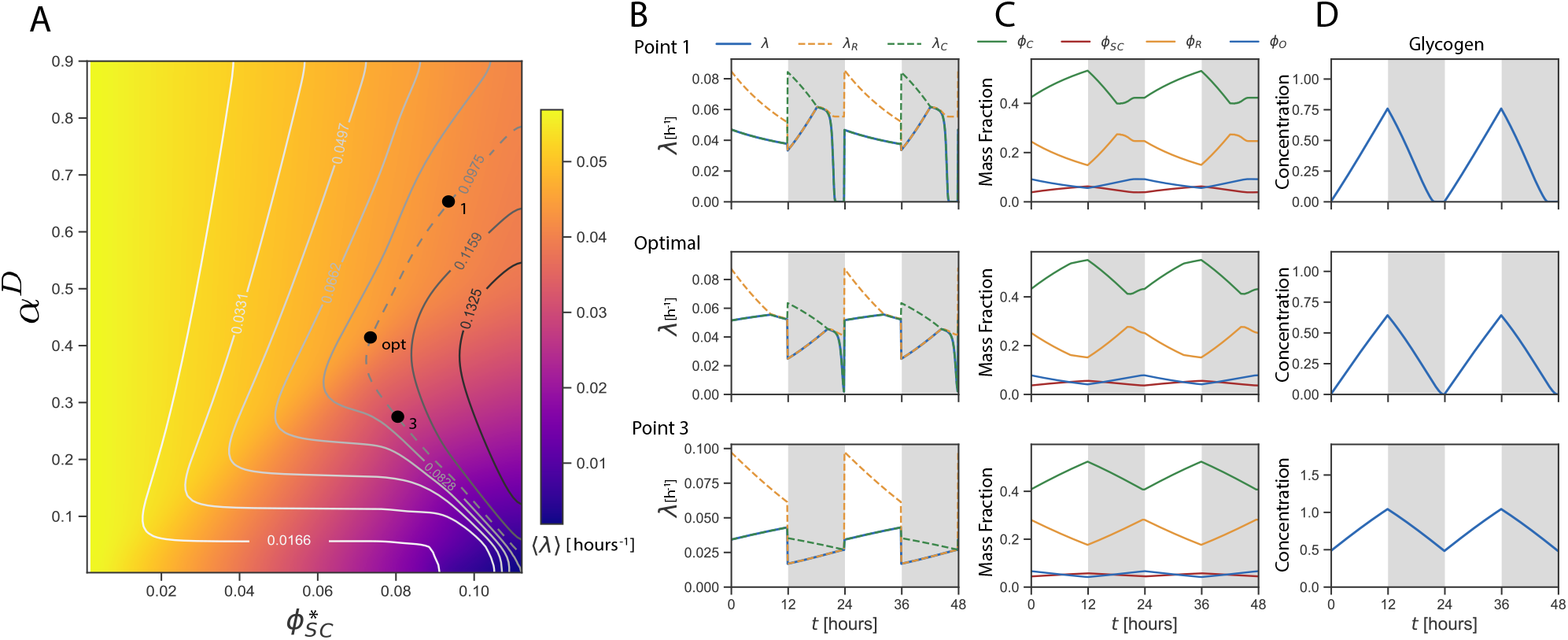
The slow relaxation of the protein sectors impedes the implementation of the optimal growth strategy suggested by Fig. 2. (A) Heatmap of the average growth rate over 24 hrs as a function of the active fraction *α*^D^ of the carbon and ribosomal sectors at night (see Eqs. 5 and 6) and the target fraction of the glycogen-storing enzyme sector 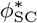 during the day. Projected onto the heatmap are contourlines of constant flux *η* through the OPPP (see Eq. 8). It is seen that along these contour lines there is an optimal, non-zero, value of *α*^D^ that maximizes the average growth rate over 24 hrs. The origin of the optimal value of *α*^D^ is elucidated in panels B-D, which show the instantaneous growth rate *λ*(*t*) (B), the protein sectors (C), and the glycogen levels *g*(*t*) (D) as a function of time, for three different points along the curve of constant *η* = 0.0975 in A. It is seen that the protein sectors relax slowly, in contrast to the dynamics in the quasi-equilibrium model (Fig. 2), in which the proteome adjusts instantly to changing nutrient levels. Moreover, even in the optimal configuration the cells grow at night (*α*^D^ *>* 0), while in the quasi-equilibrium model they do not (*α*^D^ *→* 0). In the optimal configuration, the average growth rate is significantly lower in the slow-proteome model, ⟨*λ*⟩ = 0.044*/*h, than in the quasi-equilibrium model, ⟨*λ*⟩ = 0.053*/*h (Fig. 2A). For the system marked by point 1, the active fraction *α*^D^ and hence the growth rate is relatively high during the night. The high growth rate requires a high glycogen level at the beginning of the night, but also causes this level to fall rapidly during the night. Crucially, the higher growth rate at night also means that more carbon needs to be stored during the day, which lowers the growth rate during the day. When the active fraction *α*^D^ and hence the growth rate at night is however too low (point 3), the OPP sector *ϕ*_O_ cannot rise fast enough at night, such that a very large glycogen level is required at night to meet the flux constraint *η*^*\**^. The dynamics of the slow-proteome model is fully specified by Eqs. 22-26, together with Eq. 16. Parameter values in *Supporting Information*.

To elucidate Fig. 3A, panels B-D show the instantaneous growth rate, the protein sectors, and the glycogen levels, for three points along the contourline of *η* = 0.0975 (the same value of *η* highlighted in Fig. 2A). It is immediately evident that, in each case, *ϕ*_O_ drops through dilution during the day and then increases at night. This is the fundamental reason that nighttime growth must occur in the slow relaxation model: There must be some *ϕ*_O_ to produce needed NAPDH and meet the flux constraint (Eq. 8), but without anticipation this *ϕ*_O_ can only begin to be produced at night, and nighttime production requires nighttime growth (Eq. 22).

Comparing system 1 to the optimal system that maximizes ⟨*λ*⟩ (Fig. 3A), one can see that system 1 has higher *α*^D^ and so grows faster at night, whereas the optimal system grows faster during the day. There are three distinct reasons for this behavior. The first two are already encountered for the quasi-equilibrium model (Fig. 2): First, because the optimal system grows more slowly during the night, less glycogen needs to be stored during the day. More carbon is thus immediately used for growth, which makes it possible to capitalize on the power of exponential growth, as explained above. Second, because less glycogen needs to be stored, less glycogen storing enzyme needs to be expressed during the day, and *ϕ*_SC_ is lower. Third, because, as explained above, in the optimal system the *ϕ*_O_ level is lower near the end of the night, it is also lower during the day. Together, the lower *ϕ*_SC_ and *ϕ*_O_ levels allow the *ϕ*_C_ sector to rise more rapidly at the beginning of the day in the optimal system. Indeed, while in the optimal system at the beginning of the day *λ*_C_ rises and manages to catch up with *λ*_R_, such that the system reaches its optimal configuration in which the two sectors are equally limiting, *λ* = *λ*_C_ = *λ*_R_, in the suboptimal system 1 *λ*_C_ goes down because *ϕ*_C_ does not rise fast enough, and never reaches *λ*_R_ (Fig. 3B).

Similarly, comparing system 3 to the optimal system shows that lowering *α*^D^ dramatically reduces the average growth rate over 24 hrs, in sharp constrast to the scenario in the quasi-equilibrium model, where the optimal value of *α*^D^ is 0 (Fig. 2A). In the slow-proteome model, a low growth rate at night, caused by the reduction of *α*^D^, slows down the relaxation of all protein sectors; *ϕ*_O_ therefore rises less rapidly. To meet the OPPP flux constraint during the night, the system must then crank up the glycogen level. As a result, in contrast to the other two examples, system 3 does not consume all of its glycogen at night. The remaining glycogen is then diluted out as growth occurs during the day and so is effectively lost; glycogen that is left over at the end of the night behaves as an unused carbon sink, like the inert protein sector *ϕ*_Q_.

Clearly, slow relaxation of the proteome hampers growth. Within the slow relaxation model, the cell cannot begin producing the *ϕ*_O_ it needs at night until night has actually fallen. The resulting nighttime protein production requires nighttime growth, which we know (Fig. 2) is inefficient. In fact, Fig. 3 suggests that growing at night is so costly that an alternative strategy, which can also be implemented without anticipation, might be preferable: Rather than expressing OPPP proteins only at night, the cell could instead express them at the same, constant level throughout the day and night. At nightfall, it would then already have the proteins it needed for nighttime metabolism, eliminating the necessity of night-time growth and limiting the amount of carbon that had to be stored as glycogen. This benefit would come at the cost, however, of diverting carbon early in the day from growth-promoting processes into the synthesis of *ϕ*_O_ proteins. In the *Supporting Information*, we show that, for the optimal value of *ϕ*_O_, this tradeoff is favorable: the *constant ϕ*_O_ *model* yields a somewhat higher growth rate than the slow-proteome model. In the next section, however, we will see that anticipation makes it possible to avoid the detrimental effects of slow proteome relaxation by mounting a response ahead of time, leading to a higher growth rate than in either anticipation-free scenario.

### C. Anticipation model

The anticipation model differs from the slow-proteome model primarily in that the cell stops expressing SC sector proteins and starts expressing O sector proteins a time *T*_*a*_—the anticipation time—before nightfall. To understand how much anticipation can help growth, we optimized the average growth rate over the nighttime active fraction *α*^D^ and the daytime target carbon-storing sector 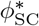 for each value of *T*_a_, and for the same values of the other parameters as in the slow-proteome model. Fig. 4 shows the result. Switching from SC to O sector expression ~3.5h before nightfall can speed up growth by about 10% compared to the slow-proteome model and by several percent compared to the constant-*ϕ*_O_ model (Fig. 4A; *Supporting Information*). Moreover, panel B shows that, for this value of the anticipation time *T*_a_, the value of *α*^D^ that maximizes the growth rate is close to zero, implying that the cells don’t grow at night. Panel C shows that this optimal configuration also corresponds to the smallest glycogen storing fraction. The benefit of anticipation can equivalently be expressed by calculating how much it allows *ϕ*_R,max_ to be reduced without lowering the growth rate below its value without anticipation—that is, how much additional protein expression that doesn’t contribute directly to growth anticipation enables. Fig. S7 in the *Supporting Information* shows that *ϕ*_R,max_ can be decreased by about 0.06 before the growth rate in the anticipation model reaches that of the slow-proteome model; the equivalent figure for the constant *ϕ*_O_ model is about 0.02. Importantly, both figures are substantially higher than the mass fraction *ϕ*_Kai_ *~* 0.002 of the core Kai clock proteins in *S. elongatus* [7].

**FIG. 4:**
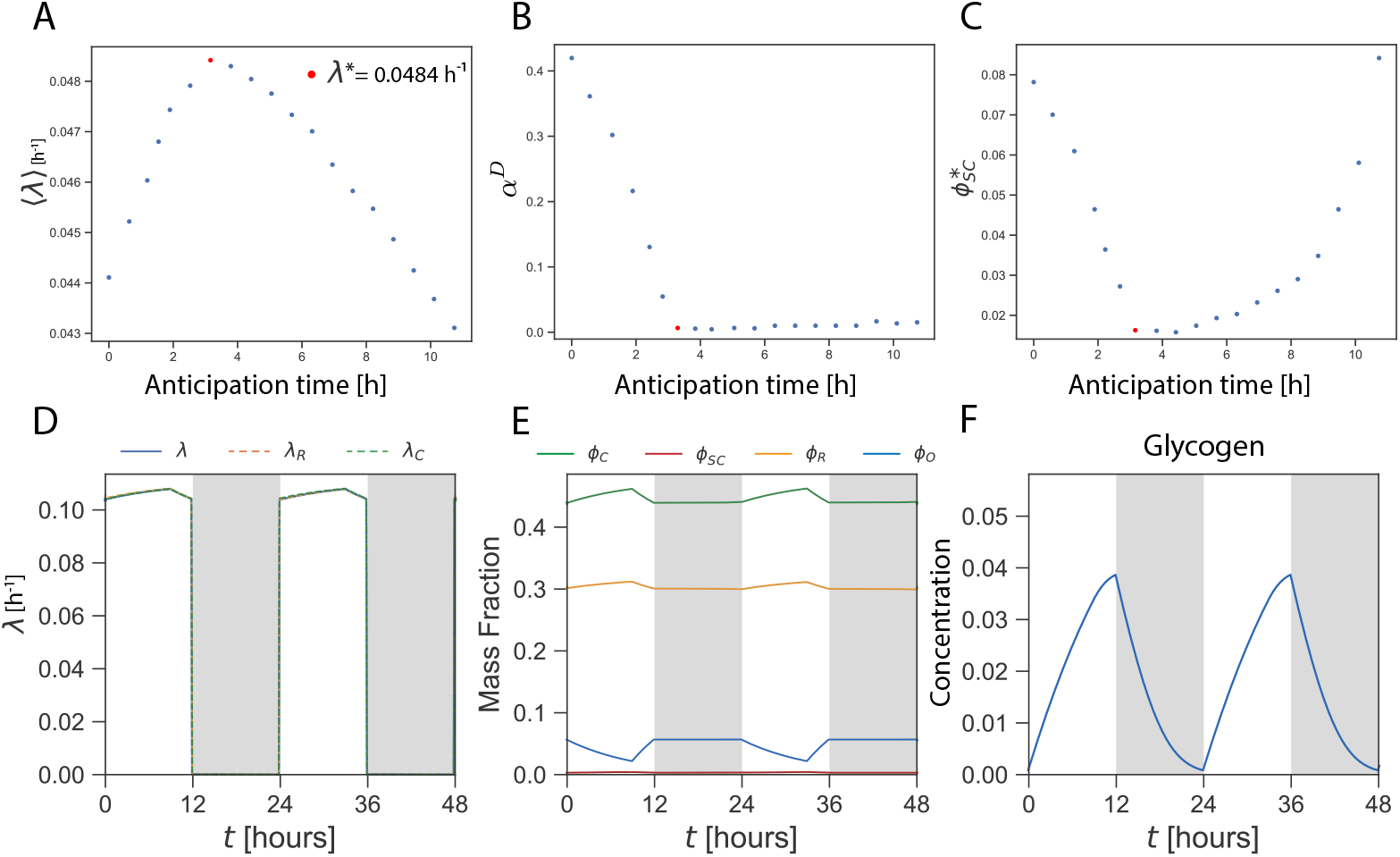
Anticipation increases the growth rate by expressing the OPPP sector at the end of the day, making it possible to shut off growth at night. Shown are results for the anticipation model, where we have optimized over *α*^D^ and 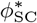 to maximize the growth rate. (A) The average growth rate over 24 hours, ⟨*λ*⟩, as a function of the anticipation time *T*_a_. There is an optimal anticipation time 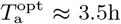 that maximizes the average growth rate. This maximal growth rate is about 10% higher than in the slow-proteome model, corresponding to *T*_a_ = 0 (see Fig. 3A). (B) The nighttime activity fraction *α*^D^ of the C and R sectors. Anticipation decreases the optimal value of *α*^D^ that maximizes the growth rate. In the optimal system, corresponding to *T*_a_ *≈* 3.5h (panel A), *α*^D^ *≈* 0, which means that the cells don’t grow at night (panel D). (C) The glycogen-storing enzyme sector *ϕ*_SC_ as a function of the anticipation time. It is seen that at the optimal anticipation time *T*_a_ = 3.5h, *ϕ*_SC_ is minimized. (D-F) Time traces of the optimal system that maximizes the growth rate, with *T*_a_ = 3.5h (see panel A): growth rate *λ*(*t*) (D), protein sectors (E), glycogen levels *g*(*t*) (F). Note that the growth rate during the day first rises because *ϕ*_O_ falls, allowing *ϕ*_R_ and *ϕ*_C_ and hence *λ* to rise. However, *T*_a_ = 3.5h before the beginning of the night, the growth rate goes down, because the cell prepares for the night by expressing the OPPP sector *ϕ*_O_ (E). Before the end of the day, *ϕ*_O_ has reached the required level for the night. Concomitantly, the growth rate is now zero during the night, in contrast to the scenario in the slow-proteome model where *ϕ*_O_ has to be made during the night, and the cells therefore need to grow during the night, see Fig. 3. The anticipation model is identical to the slow-proteome model given by Eqs. 22-26, together with Eq. 16, except that *ϕ*_SC_ expression stops and *ϕ*_O_ expression starts an anticipation time *T*_a_ before nightfall, and the reverse happens *T*_a_ before daybreak. Parameter values in *Supporting Information*.

To elucidate this behavior, we show in Fig. 4D-F time traces at the optimal anticipation time 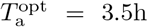 (Fig. 4A). As in the slow-proteome model, the growth rate (panel D) rises early in the day as the *ϕ*_O_ level falls (panel E). This trend reverses when the anticipation time is reached and O sector proteins begin to be expressed. Clearly, there is a cost to anticipation: it can lower the instantaneous growth rate. The cell should therefore not express *ϕ*_O_ proteins too early in the day. Yet, expressing these enzymes during the day also has a marked benefit: it makes it possible to reach a high enough *ϕ*_O_ level before nightfall, eliminating the need for *ϕ*_O_ expression and the associated growth at night. As a result, only enough glycogen has to be stored to meet the OPPP flux constraint, with no additional glycogen needed to fuel nighttime growth. The levels of glycogen are indeed dramatically lower than in the slow-proteome model (compare Fig. 4F to Fig. 3D). As discussed above, the reduction in the amount of carbon that needs to be stored has two important consequences: the carbon-storing sector *ϕ*_SC_ can be smaller, so that more proteins can be made that directly contribute to growth, and more carbons can be used to fuel growth immediately, taking advantage of exponential growth. In summary, anticipation makes it possible to implement the optimal growth strategy suggested by the quasi-equilibrium model (see Fig. 2), namely to grow exclusively during the day, even when the proteome relaxes slowly, on a timescale set by *λ*.

## III. ESTIMATING PROTEIN MASS REDISTRIBUTION IN *S. ELONGATUS* FROM PROTEOMIC DATA

A central premise of our model is that the cells must repartition their proteome among the different sectors over the course of 24 hrs in order to meet their different physiological needs in light and dark environments, leading to significant changes in the protein mass distribution across the different sectors (Fig. 4E). Here, we analyse the proteomics data of Ref. [7], obtained under Light-Dark (LD) cycles, to determine whether the cyanobacterium *S. elongatus* does indeed appreciably redistribute its protein mass across the different sectors over the course of 24 hrs and whether this redistribution is consistent with our model.

### A. The PMHA Method

To analyse the mass redistribution, we present a new technique called Principal Mass Harmonics Analysis (PMHA), which is based on approximating the measured protein levels by the first harmonics of their 24-hour oscillations (Fig. 5A). Existing data reduction approaches for similar proteomics data often try to identify a subset of individual proteins with a high circadian oscillation amplitude or a specific oscillation phase. PMHA instead seeks to determine globally whether there is a significant circadian redistribution of mass across the proteome. We show that this question can be answered with confidence even when the changes in the mass distribution are at the ~10% level, well below the oscillation amplitude that would be reliably detectable for a single protein in typical mass spectrometry data.

**FIG. 5:**
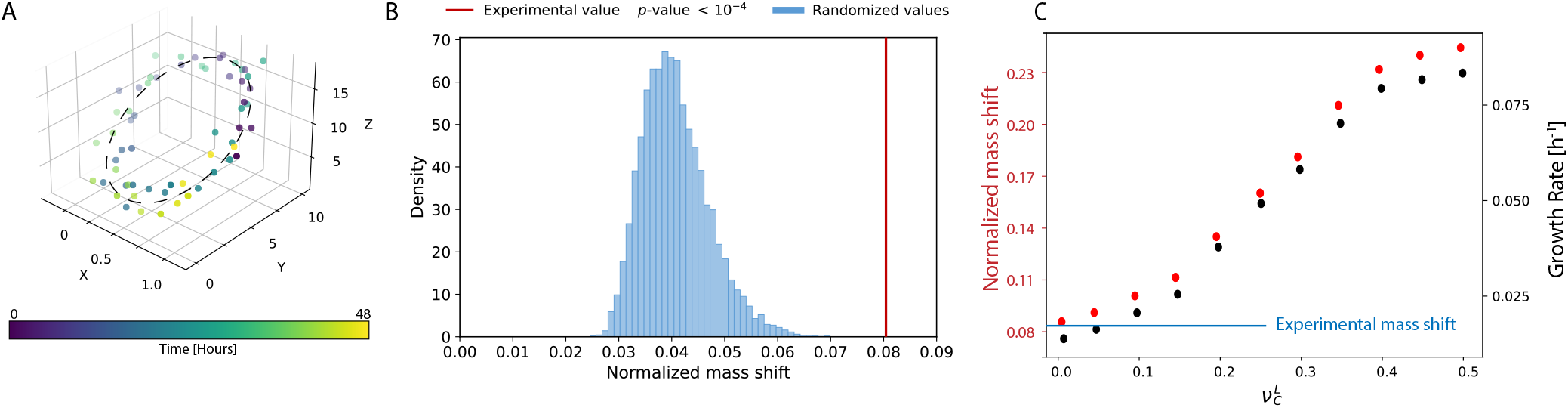
The PMHA method. A) Schematic of the method for the case where only *N* = 3 protein levels are measured, corresponding to the *X, Y*, and *Z* axes. The points represent the measured levels of these proteins at different times (color scale below). The PMHA method approximates the cycle traced out by these noisy data points as an ellipse (dashed curve) and takes the major axis of this ellipse, normalized by the distance from the origin to the center of the ellipse, as a measure of how much the proteome mass redistributes over 24 hours, i.e. of the mass shift. B) Histogram showing the density (defined so that the histogram integrates to 1) of normalized mass shift values obtained from 10,000 randomizations of the experimental time traces. The red line indicates the actual mass shift computed from the mass spectrometry data. C) Normalized mass shift and growth rate as functions of 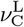 for our model. The experimental mass shift obtained by applying PMHA to the data of [7] is indicated in blue, corresponding to a model system receiving relatively low intensity illumination with a cell division time of approximately 48 hours.

Specifically, after appropriate transformation and normalization (see *Supporting Information*), the data of [7] can be viewed as providing the mass fraction *X*_*i*_(*t*) as a function of time *t* for each protein *i* in the data set. (We distinguish the mass fraction *X*_*i*_ of an individual protein from the mass fraction *ϕ*_*α*_ of an entire sector.) We approximate *X*_*i*_(*t*) as

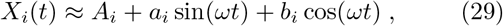

where *ω* = 2*π/*(24h) is the fundamental frequency of the daily oscillation and *A*_*i*_, *a*_*i*_, and *b*_*i*_ are constants; we thus ignore all higher harmonics with frequencies that are multiples of *ω*.

The vector with components *X*_*i*_(*t*) given by Eq. 29 traces out an ellipse as a function of time in an *N*-dimensional space, as represented schematically in Fig. 5A for *N* = 3 different protein species. In effect, we approximate the noisy experimental trajectory as an ellipse. The major axis of this ellipse serves as a measure of how much the proteome composition changes over one day-night cycle. Specifically, we use as our figure of merit the “normalized mass shift”, which we define as the length of this major axis normalized by the distance 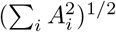 from the origin to the center of the ellipse.

To evaluate the significance of this normalized mass shift, we compare the observed shift with a randomized control. This control is generated by applying the same procedure to shuffled versions of the original time traces *X*_*i*_(*t*). By repeating this randomization multiple times, we construct a histogram of the resulting mass shifts and compute a *p*-value by measuring the fraction of random outcomes that exceed the observed shift. This comparison allows us to assess whether the detected mass redistribution is significantly different from random fluctuations.

### B. PMHA Results

Fig. 5B presents the results of our PMHA analysis applied to the *S. elongatus* data of [7]. We find that the normalized mass shift is slightly more than 0.08, indicating that about 8% of the total proteome mass is redistributed between day and night. This shift is highly statistically significant; in fact, none of our randomized data sets had shifts this large, allowing us to estimate a *p*-value *p≤* 1.7 *×* 10^*−*6^.

To compare these experimental results to the predictions of our theoretical framework, we performed exactly the same PMH analysis on the time traces of the protein sectors of our anticipation model (Fig. 4E) for different values of the parameter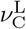, corresponding to different light intensities and hence to different growth rates. For each 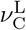, we maximized the growth rate as before by optimizing over the target carbon-storing sector fraction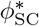, the maximum activity *α*^D^ of the C and R sectors at night, and the anticipation time *T*_a_.

Fig. 5C shows the result. As 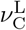 increases, the growth rate rises and then saturates. The plateau level is set by the maximum translation rate *ν*_R_, the maximum size of the ribosomal sector *ϕ*_R,max_, and the requirement that both *ϕ*_SC_ and *ϕ*_O_ must be large enough to meet the OPPP flux constraint at night (see Eqs. 8 and 9).

Fig. 5C also shows that the normalized mass shift increases with the growth rate, reaching nearly 25% at the highest growth rates (at the limit of what is attainable in the standard *S. elongatus* PCC 7942 strain in LD 12:12 [39, 49]). This behavior is readily understood from the fact that the growth rate *λ* sets the timescale for changes in proteome composition: at higher growth rates, larger changes in the proteome can be achieved during a given 12h day or night period. In panel C, we have also marked the *~* 8% normalized mass shift from the experimental data of [7]. This lower figure suggests that the growth rate of the cells under the conditions of [7] is *λ*_exp_ *≈* 0.015h^*−*1^, corresponding to a cell division time of close to 48 hrs. This is consistent with the relatively low light levels of 80*µ*M photons m^*−*2^s^*−*1^ reported in [7] (see also [40]).

## IV. DISCUSSION

It is an intriguing observation that most cyanobacteria grow exclusively during the day. After all, these cells can store carbon during the day, which could, in principle, be used to fuel growth at night. To understand their circadian metabolism, we developed a model based on the framework of Hwa and coworkers [22, 23, 27, 28, 31].

The power of this framework is that it provides a coarse-grained description of the proteome with only a limited number of sectors, characterized by enzyme efficiencies that can be measured experimentally [22, 23, 27, 28, 31]. Our model is arguably the simplest than can capture the tradeoffs between using carbon to drive growth during the day and storing carbon to fuel processes at night. It consists of a ribosome (R) sector, a carbon (C) sector that fixes the carbon for protein synthesis, a sector (SC) that stores carbon during the day to fuel essential processes at night, and a sector (O) that catalyzes these nighttime processes, consuming the stored carbon (Fig. 1). While our model is minimal, its dynamics is highly non-linear. In particular, it reveals that the optimal strategy that maximizes the average growth rate over 24 hrs is not to grow at night at all. Our analysis identifies two fundamental reasons why this strategy is optimal: 1) to capitalize on the power of exponential growth, it is beneficial to be greedy, and thus to use building blocks for growth as soon as they become available, so that they can in turn fix more carbon and synthesize more protein; and storing building blocks requires enzymes (the SC sector), which use resources that then cannot be devoted to fueling growth directly. Consequently, the optimal strategy is to store only enough carbon as glycogen to fuel processes that are essential for nighttime survival.

Our analysis also reveals a severe challenge in implementing this optimal strategy. In the absence of active protein degradation, cells must grow in order to adjust their proteome. Yet, cyanobacterial cells grow slowly, which means that the timescales to adjust the proteome through growth will be comparable to or longer than the 24h period of the day-night rhythm. As a result, in the absence of anticipation, the cells that do not express *ϕ*_O_ during the day would need to grow significantly during the night in order to reach the necessary size for the OPPP sector *ϕ*_O_. Indeed, the principal benefit of having a circadian clock, according to our model, is that it allows the cyanobacterium to anticipate the shift from day to night and hence to express the *ϕ*_O_ sector only late in the day; in this way, the cell avoids both the costs associated with nighttime growth and those associated with expressing *ϕ*_O_ throughout the day when it is not needed. Without anticipation, the cell cannot implement this strategy. The predicted anticipatory expression appears to be consistent with transcriptomic and proteomic data. [2, 4, 5, 7].

Our model’s central prediction concerns the benefit of clock-enabled anticipation: Fig. 4A shows that the average growth rate at the optimal anticipation time *T*_a_ *≈* 3.5h is about 10% higher than the average growth rate without anticipation (*T*_a_ = 0) and about 5% higher than the growth rate for constant *ϕ*_O_ expression (*Supporting Information*). Both figures substantially outstrip the cost of expressing the KaiABC clock proteins (*Supporting Information* Fig. S7). Thus, our work both explains why clock-enabled anticipation is beneficial and demonstrates quantitatively how it can be a powerful driver of increased fitness in a simple bacterial system.

It is worth pointing out that we have arrived at these results using the simplest possible model of anticipation, in which information from the clock is used only to implement a single, step-like switch in ribosome allocation at a single anticipation time *T*_a_. In reality, the cell can likely use different anticipation times for different parameters and vary these parameters smoothly throughout the day, rather than in a single sharp switch. Optimization over this much higher dimensional parameter space might lead to further gains in growth rate. For example, recent evidence indicates that *S. elongatus* limits glycogen synthesis early in the day by driving condensation of glycogen synthases [50], consistent with the assumptions of earlier models [51]. By freeing more carbon for protein synthesis and exponential growth early in the day, this strategy would be expected to increase the average growth rate through essentially the same mechanism identified by our quasi-equilibrium model, namely that using carbon for immediate growth, which in turn increases carbon fixation capacity, is more efficient than saving it for later.

The formalism we have developed makes additional predictions that can be tested experimentally. For example, Fig. 5C gives the growth rate and normalized proteome mass shift as a function of the daytime carbon sector efficiency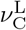, which depends on the light intensity and carbon dioxide concentration; in the simplest case where carbon dioxide is in excess, 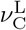 is expected to be proportional to the light intensity. Similarly, in the *Supporting Information* (Fig. S7) we calculate the maximum growth rate as a function of the parameter *ϕ*_R,max_, which can be lowered experimentally by ectopic expression of an inert protein [27]. A beautiful series of experiments show that cyanobacteria grow fastest when their clock period is matched to the period of the light-dark (LD) cycle imposed on them [17, 18]; the likely reason for slower growth when the period is mismatched is that the clock entrains to the LD cycle with the wrong phase—or, in the language of this paper, with the wrong *T*_a_. Fig. 4A predicts the growth rate as a function of *T*_a_. We note that in all of these examples, we assume that the cell can optimize over 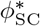 and *α*^D^ for each new value of the parameter varied by the experimentalist; if this is not the case, growth rates will be lower than predicted. More accurate predictions would require some data on how these various parameters actually depend on each other for a given experimental protocol. One promising first step might be to measure the time course of glycogen storage in different conditions; together with measured growth rates, this can in principle be used to infer relative values of 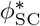 and also to determine the net nighttime flux *η* through the OPPP (Eq. 8).

While the dynamics of our minimal model is already complex, it may be useful to add sectors as more data becomes available. In particular, it might be of interest to distinguish between proteins of a given sector that are generic, i.e. expressed at significant levels both during the day and during the night, and proteins that are specific to one part of the day, such as the photosystem components. The challenge will be to define major subsectors and devise experiments which make it possible to measure the associated enzyme efficiences.

The model that we have introduced describes cyanobacteria like *S. elongatus* and *Synechocystis* that fix carbon during the day. However, the framework can also be extended to the metabolism of cyanobacteria such as *Cyanothece* [52], which fix not only carbon during the day but also nitrogen during the night. Interestingly, this bacterium also adopts an extreme growth strategy, in which it grows exclusively during the day.

## Acknowledgments

We are grateful to Maarten Altelaar, Ilka Axmann, and Albert Heck for sharing the data published in [7] and for helping with its intepretation. This work was funded in part by NSF grant DMR-2243624 to DKL. PRtW performed his work at the research institute AMOLF and acknowledges support from the Dutch Research Council (NWO) and from the European Research Council (ERC) under the European Union’s Horizon 2020 research and innovation program (grant agreement 885065).

MM, DKL and PRtW wrote the manuscript; MM performed research and analyzed data; MM, DKL and PRtW designed research. The authors declare that they have no conflicts or competing interests.

## References

[1] Pattanayak, G. K., Phong, C. & Rust, M. J. Rhythms in Energy Storage Control the Ability of the Cyanobacterial Circadian Clock to Reset. Current Biology 24, 1934–1938 (2014).

[2] Diamond, S., Jun, D., Rubin, B. E. & Golden, S. S. The circadian oscillator in Synechococcus elongatuscontrols metabolite partitioning during diurnal growth. Proc Natl Acad Sci U S A 112, E1916–E1925 (2015).

[3] Liu, Y. et al. Circadian orchestration of gene expression in cyanobacteria. Genes & Development 9, 1469–1478 (1995).

[4] Ito, H. et al. Cyanobacterial daily life with Kai-based circadian and diurnal genome-wide transcriptional control in Synechococcus elongatus. Proceedings of the National Academy of Sciences 106, 14168–14173 (2009).

[5] Vijayan, V., Zuzow, R. & O’Shea, E. K. Oscillations in supercoiling drive circadian gene expression in cyanobacteria. Proc Natl Acad Sci U S A 106, 22564–22568 (2009).

[6] Markson, J. S., Piechura, J. R., Puszynska, A. M. & O’Shea, E. K. Circadian Control of Global Gene Expression by the Cyanobacterial Master Regulator RpaA. Cell 155, 1396–1408 (2013).

[7] Guerreiro, A. C. L. et al. Daily rhythms in the cyanobacterium synechococcus elongatus probed by high-resolution mass spectrometry-based proteomics reveals a small defined set of cyclic proteins. Molecular & Cellular Proteomics 13, 2042–2055 (2014). URL https://www.ncbi.nlm.nih.gov/pmc/articles/PMC4125736/.

[8] Diamond, S. et al. Redox crisis underlies conditional light-dark lethality in cyanobacterial mutants that lack the circadian regulator, RpaA. Proc Natl Acad Sci U S A 114, E580–E589 (2017).

[9] Latifi, A., Ruiz, M. & Zhang, C.-C. Oxidative stress in cyanobacteria. FEMS Microbiology Reviews 33, 258–278 (2009).

[10] Shinde, S. et al. Glycogen metabolism supports photosynthesis start through the oxidative pentose phosphate pathway in cyanobacteria. Plant Physiology 182, 507–517 (2020). URL 10.1104/pp.19.01184.

[11] Liao, Y. & Rust, M. J. The circadian clock ensures successful dna replication in cyanobacteria. Proc Natl Acad Sci U S A 118, e2022516118 (2021).

[12] Welkie, D. G. et al. Genome-wide fitness assessment during diurnal growth reveals an expanded role of the cyanobacterial circadian clock protein KaiA. Proc Natl Acad Sci U S A 115, E7174–E7183 (2018).

[13] Puszynska, A. M. & O’Shea, E. K. Switching of metabolic programs in response to light availability is an essential function of the cyanobacterial circadian output pathway. eLife 6, e23210 (2017).

[14] Reimers, A.-M., Knoop, H., Bockmayr, A. & Steuer, R. Cellular trade-offs and optimal resource allocation during cyanobacterial diurnal growth. Proc Natl Acad Sci U S A 114, E6457–E6465 (2017).

[15] Johnson, C. G. M. et al. Multi-omics reveals temporal scales of carbon metabolism in synechococcus elongatus pcc 7942 under light disturbance. PRX Life 3, 033017 (2025). URL https://link.aps.org/doi/10.1103/l2dp-kw2t.

[16] Johnson, C. G. M. et al. Spatiotemporal 4d wholecell modeling of a minimal autotroph reveals central carbon metabolism regulated locally by protein megacomplexes via post-translational modifications under light disturbance. bioRxiv 2026.01.21.700212 (2026). URL https://www.biorxiv.org/content/early/2026/01/23/2026.01.21.700212.

[17] Ouyang, Y., Andersson, C. R., Kondo, T., Golden, S. S. & Johnson, C. H. Resonating circadian clocks enhance fitness in cyanobacteria. Proc Natl Acad Sci U S A 95, 8660–8664 (1998).

[18] Woelfle, M. A., Ouyang, Y., Phanvijhitsiri, K. & Johnson, C. H. The Adaptive Value of Circadian ClocksAn Experimental Assessment in Cyanobacteria. Current Biology 14, 1481–1486 (2004).

[19] Becker, N. B., Mugler, A. & ten Wolde, P. R. Optimal Prediction by Cellular Signaling Networks. Physical Review Letters 115, 258103 (2015).

[20] Tjalma, A. J. et al. Trade-offs between cost and information in cellular prediction. Proceedings of the National Academy of Sciences 120, e2303078120 (2023).

[21] Van den Ende, M. Circadian Metabolism: Modeling the Metabolic States of Circadian Cells. Master’s thesis, Universiteit van Amsterdam (2017).

[22] Erickson, D. W. et al. A global resource allocation strategy governsgrowth transition kinetics of Escherichia coli. Nature 551, 119–123 (2017).

[23] Mori, M., Schink, S., Erickson, D. W., Gerland, U. & Hwa, T. Quantifying the benefit of a proteome reserve in fluctuating environments. Nature Communications 8, 1–7 (2017).

[24] Sinetova, M. A., Červený, J., Zavňel, T. & Nedbal, L. Journal of Biotechnology. Journal of Biotechnology 162, 148–155 (2012).

[25] Teng, S. W., Mukherji, S., Moffitt, J. R., de Buyl, S. & O’Shea, E. K. Robust Circadian Oscillations in Growing Cyanobacteria Require Transcriptional Feedback. Science 340, 737–740 (2013).

[26] Bernstein, H. C. et al. Effect of mono- and dichromatic light quality on growth rates and photosynthetic performance of Synechococcus sp. PCC 7002. Frontiers in Microbiology 5, 488 (2014).

[27] Scott, M., Gunderson, C. W., Mateescu, E. M., Zhang, Z. & Hwa, T. Interdependence of Cell Growth and Gene Expression: Origins and Consequences. Science 330, 1099– 1102 (2010).

[28] You, C. et al. Coordination of bacterial proteome with metabolism by cyclic AMP signalling. Nature 500, 1–6 (2013).

[29] Deris, J. B. et al. The Innate Growth Bistability and Fitness Landscapes of Antibiotic-Resistant Bacteria. Science 342, 1237435–1237435 (2013).

[30] Li, G.-W., Burkhardt, D., Gross, C. & Weissman, J. S. Quantifying Absolute Protein Synthesis Rates Reveals Principles Underlying Allocation of Cellular Resources. Cell 157, 624–635 (2014).

[31] Hui, S. et al. Quantitative proteomic analysis reveals a simple strategy of global resource allocation in bacteria. Molecular Systems Biology 11, e784–e784 (2015).

[32] Monti, M., Lubensky, D. K. & ten Wolde, P. R. Robustness of clocks to input noise. Physical Review Letters 121, 078101 (2018).

[33] Pittayakanchit, W., Lu, Z., Chew, J., Rust, M. J. & Murugan, A. Biophysical clocks face a trade-off between internal and external noise resistance. eLife 7, e37624 (2018).

[34] Schober, A. F., Holmes, C., Palmer, S. & Rust, M. J. The free-running property of circadian clocks is needed to tolerate changing photoperiods. Current Biology 36, 1548–1555.e5 (2026).

[35] Dani, C. et al. Seasonal cycles select for self-sustained circadian oscillators. Current Biology 36, 1541–1547 (2026).

[36] Stöckel, J. et al. Diurnal Rhythms Result in Significant Changes in the Cellular Protein Complement in the Cyanobacterium Cyanothece 51142. PLoS ONE 6, e16680 (2011).

[37] Aryal, U. K. et al. Dynamic proteomic profiling of a unicellular cyanobacterium Cyanothece ATCC51142 acrosslight-dark diurnal cycles. BMC Systems Biology 5, 194 (2011).

[38] Zwicker, D., Lubensky, D. K. & ten Wolde, P. R. Robust circadian clocks from coupled protein-modification and transcription-translation cycles. Proc Natl Acad Sci U S A 107, 22540–5 (2010).

[39] Ungerer, J., Lin, P.-C.Chen, H.-Y. & Pakrasi, H. B. Adjustments to Photosystem Stoichiometry and Electron Transfer Proteins Are Key to the Remarkably Fast Growth of the Cyanobacterium Synechococcus elongatus UTEX 2973. mBio 9, e02327–17 (2018).

[40] Broddrick, J. T. et al. Predicting the metabolic capabilities of Synechococcus elongatus PCC 7942 adapted to different light regimes. Metabolic Engineering 52, 42–56 (2019).

[41] Pattanayak, G. K. et al. Daily Cycles of Reversible Protein Condensation in Cyanobacteria. Cell Reports 32, 108032 (2020).

[42] Dennis, P. P. & Bremer, H. Macromolecular composition during steady-state growth of Escherichia coli B-r. Journal of Bacteriology 119, 270–281 (1974).

[43] Goldberg, A. L. & St. John, A.C. Intracellular protein degradation in mammalian and bacterial cells: Part 2. Annual Review of Biochemistry 45, 747–804 (1976). URL https://www.annualreviews.org/content/journals/10.1146/annurev.bi.45.070176.003531.

[44] Chen, W. et al. A systematic survey of the light/dark-dependent protein degradation events in a model cyanobacterium. Molecular & Cellular Proteomics 20 (2021). URL https://doi.org/10.1016/j.mcpro.2021.100162. Doi: 10.1016/j.mcpro.2021.100162.

[45] Kolter, R., Balaban, N. & Julou, T. Bacteria grow swiftly and live thriftily. Current Biology 32, R599– R605 (2022). URL https://doi.org/10.1016/j.cub.2022.05.005. Doi: 10.1016/j.cub.2022.05.005.

[46] Planqué, R., Hulshof, J., Teusink, B., Hendriks, J. C. & Bruggeman, F. J. Maintaining maximal metabolic flux by gene expression control. PLoS Computational Biology 14, e1006412 (2018).

[47] Zavňel, T. et al. Quantitative insights into the cyanobacterial cell economy. eLife 8, e42508 (2019).

[48] Hood, R. D., Higgins, S. A., Flamholz, A., Nichols, R. J. & Savage, D. F. The stringent response regulates adaptation to darkness in the cyanobacterium synechococcus elongatus. Proceedings of the National Academy of Sciences 113, E4867–E4876 (2016).

[49] Ungerer, J., Wendt, K. E., Hendry, J. I., Maranas, C. D. & Pakrasi, H. B. Comparative genomics reveals the molecular determinants of rapid growth of the cyanobacterium Synechococcus elongatus UTEX 2973. Proceedings of the National Academy of Sciences 115, E11761– E11770 (2018).

[50] Burton, L. A., Pattanayak, G. K. & Rust, M. J. Rhythmic protein condensation in the circadian regulation of glycogen metabolism. iScience 29, 115605 (2026).

[51] Hellweger, F. L., Jabbur, M. L., Johnson, C. H., van Sebille, E. & Sasaki, H. Circadian clock helps cyanobacteria manage energy in coastal and high latitude ocean. The ISME Journal 14, 560–568 (2020). URL 10.1038/s41396-019-0547-0.

[52] Monti, M., Lubensky, D. K. & ten Wolde, P. R. Theory of circadian metabolism. arXiv 1805.04538 (2018).

