## Supplemental text and figures (1 file) for "Growth, metabolism, and the benefits of clock-enabled anticipation"

### Supporting information for "Growth, metabolism, and the benefits of clock-enabled anticipation"

**Contents**

**I. qORAC and the Slow Proteome Model** 1
A. Basic qORAC 1
B. A simple limiting case 6

**II. Dependencies on $\phi_{\rm O}$** 7
A. $\phi^*_{\rm O}$ in the anticipation model 7
B. Constant $\phi_{\rm O}$ model 7

**III. Principle Mass Harmonics Analysis (PMHA) Method** 8

**IV. Methods** 10
A. Model parameters 10
B. Numerical methods 11

**V. Supporting Figures** 11

**References** 12

In this supporting information, we first (in Section I) outline one possible mechanistic basis for the central assumptions of the slow-proteome model introduced in the main text. We then (in Section II) elaborate on how the growth rates that we calculate depend on the target mass fraction $\phi^*_{\rm O}$ for the OPP pathway and justify our choice not to exhaustively optimize over this parameter; we also discuss the constant $\phi_{\rm O}$ model, in which $\phi_{\rm O}$ is held at one, fixed level throughout the day and night. Section III provides additional technical details on the Principle Mass Harmonics Analysis (PMHA) method, introduced in the main text, for estimating the extent of proteome mass redistribution between daytime and nighttime. Section IV discusses how we estimate appropriate parameter values for the model and touches on numerical methods; it also includes a table of the parameter values used throughout the main text. Finally, we include several supporting figures related to figures in the main text.

#### I. QORAC AND THE SLOW PROTEOME MODEL

In this section, we show how several key assumptions of our slow-proteome model can be derived from a minimal version of the qORAC model originally proposed to explain aspects of the physiology of non-photosynthetic micro-organisms like *E. coli* [1]. In the main text, we argued that, in order to generalize the quasi-equilibrium model to the slow proteome model, we need to know how the cell 1) adjusts the activities of the carbon and ribosome sectors to set its growth rate and 2) chooses its ribosome allocation fractions $\chi_\alpha(t)$ away from steady state. To answer these questions, we introduced a set of straightforward assumptions (Eqs. 23 - 28 of the main text) that we justified on a heuristic, phenomenological basis. Our goal in this section is to show how these same assumptions also arise naturally as a limiting case of a simple, mechanistic model of cell physiology.

To this end, we illustrate the main ideas with a minimal version of the qORAC model [1] in which there are only two active protein sectors, carbon C and ribosome R, as well as an inactive sector Q, so that $\phi_{\rm R} + \phi_{\rm C} = \phi_{\rm R,max} = 1 - \phi_{\rm Q}$. Below, we first introduce the basic ideas of the qORAC model without taking any special limits, and then we show how the simple rules used in the main text are recovered as a limiting case of this model.

##### A. Basic qORAC

The qORAC model imagines that the carbon sector produces an amino acid pool that is in turn consumed by the ribosome sector to produce proteins. The fluxes through the carbon and ribosome sectors are modulated as a function of the level of the amino acid pool to prevent catastrophic amino acid build-up or depletion.

Specifically, let $A(t)$ be the total mass of free amino acids in this pool, and define $a(t) = A(t)/M(t)$ to be the amino acid mass normalized by the total protein mass $M(t)$. (Note that $M(t)$ is not defined to include the mass of free amino acids, so $a(t)$ can be compared to the protein mass fractions $\phi_\alpha$ but is not strictly itself a mass fraction. Below, we sometimes loosely use the term "concentration" for $a(t)$, even though it is normalized by mass rather than by volume.) In a simple model in which the amino acid pool is produced by the C sector and consumed to make proteins by the R sector, the amino acid pool evolves according to

$$\frac{dA}{dt} = \nu_{\rm C}(t)\alpha_{\rm C}(t)M_{\rm C}(t) - \nu_{\rm R}\alpha_{\rm R}(t)M_{\rm R}(t) \tag{S1}$$

$$\frac{da}{dt} = J_{\rm C}(t) - J_{\rm R}(t) - \lambda a(t) \tag{S2}$$

$$= \nu_{\rm C}(t)\alpha_{\rm C}(t)\phi_{\rm C}(t) - \nu_{\rm R}\alpha_{\rm R}(t)\phi_{\rm R}(t) - \lambda a(t) \tag{S3}$$

$$\approx \nu_{\rm C}(t)\alpha_{\rm C}(t)\phi_{\rm C}(t) - \nu_{\rm R}\alpha_{\rm R}(t)\phi_{\rm R}(t)\ , \tag{S4}$$

where we justify below dropping the dilution term $\lambda a$, and the expressions for the fluxes through the R and C sectors

$$J_{\rm C}(t) = \nu_{\rm C}(t)\alpha_{\rm C}(t)\phi_{\rm C}(t) \tag{S5}$$

$$J_{\rm R}(t) = \nu_{\rm R}\alpha_{\rm R}(t)\phi_{\rm R}(t) \tag{S6}$$

follow directly from the corresponding equations in the main text (Eqs. 2 and 3) in the special case that the only two active sectors are C and R. In the quasi-equilibrium model, we assumed that the active fractions of the carbon and ribosome sectors are equal to each other and take one constant value throughout the day and another throughout the night, $\alpha_{\rm R}(t) = \alpha_{\rm C}(t) = \alpha^{\beta}$, with $\beta = {\rm L}, {\rm D}$. In contrast, in the qORAC-inspired model considered here and in the slow-proteome model of the main text, we allow these factors to be time-dependent and to differ between the two sectors. Specifically, to implement the necessary feedback from the amino acid pool to the fluxes into and out of it, we assume that the carbon sector is down-regulated and the ribosome sector up-regulated when the free amino acid level $a(t)$ becomes too high, and vice-versa when it becomes too low:

$$\alpha_{\rm C}(t) = \alpha^{\beta}\frac{K_{\rm C}}{a(t) + K_{\rm C}}, \tag{S7}$$

$$\alpha_{\rm R}(t) = \alpha^{\beta}\frac{a(t)}{a(t) + K_{\rm R}}, \tag{S8}$$

where $\alpha^{\beta}$ denotes the maximum activity, taken to be $\alpha^{L} = 1$ during the day and a parameter to be optimized, $\alpha^{D}$, during the night (just as in the quasi-equilibrium model).

We further assume that $K_{\rm C}, K_{\rm R} \ll \phi_{\rm C}, \phi_{\rm R}$, so that the mass of the free amino acids is much less than the mass bound up in proteins, $a \ll \phi_{\rm C}, \phi_{\rm R}$. Then, because $\lambda, \nu_{\rm C}$, and $\nu_{\rm R}$ are all of the same order, $\lambda a$ is much smaller than the other two terms in Eq. S3 and can be dropped. Moreover, because $a$ is small, a small imbalance between the fluxes through the C and R sectors can cause rapid relative changes in $a$, so that we can assume that $a(t)$ is in quasi-equilibrium with respect to the $\phi_\alpha$'s. That is, $da/dt = 0$ in Eq. S4, and

$$\nu_{\rm C}\frac{K_{\rm C}}{a + K_{\rm C}}\phi_{\rm C} = \nu_{\rm R}\frac{a}{a + K_{\rm R}}\phi_{\rm R}\ , \tag{S9}$$

or, after rearranging,

$$\frac{a(t)}{K_{\rm C}}\left(\frac{K_{\rm C} + a(t)}{K_{\rm R} + a(t)}\right) = \frac{\nu_{\rm C}(t)\phi_{\rm C}(t)}{\nu_{\rm R}\phi_{\rm R}(t)}\ . \tag{S10}$$

This equation determines $a(t)$ at each time given the values of $\phi_{\rm C}$ and $\phi_{\rm R}$. (In particular, $a(t)$ is a monotonically increasing function of the ratio $\nu_{\rm C}(t)\phi_{\rm C}(t)/\nu_{\rm R}\phi_{\rm R}(t)$.) The equation also ensures that the fluxes through the two sectors are equal, so that we can write the growth rate as

$$\lambda(t) = \nu_{\rm R}\alpha_{\rm R}(t)\phi_{\rm R}(t) = \nu_{\rm R}\alpha^{\beta}\frac{a(t)}{K_{\rm R} + a(t)}\phi_{\rm R}(t) \tag{S11}$$

$$= \nu_{\rm C}(t)\alpha_{\rm C}(t)\phi_{\rm C}(t) = \nu_{\rm C}(t)\alpha^{\beta}\frac{K_{\rm C}}{K_{\rm C} + a(t)}\phi_{\rm C}(t). \tag{S12}$$

These equations answer the first of the two main questions that must be addressed once we move from from the quasi-equilibrium model to the slow-proteome model: how does the cell adjust the activities of the carbon and ribosome sectors, thereby setting the growth rate $\lambda(t)$? We emphasize that Eqs. S11 and S12 hold at all times, even during shifts between day and night when the protein sectors are still evolving.

We now turn our attention to the second open question: What determines the fractions $\chi_\alpha$ that describe how newly synthesized proteins are allocated to the different sectors (Eqs. 27-28 in the main text)? We begin by observing that, once we allow feedback from the amino acid pool to the active fractions $\alpha_{\rm R}$ and $\alpha_{\rm C}$, the free amino acid level will adjust so that the fluxes through the sectors are balanced, and any choice of the sector fractions $\phi_{\rm C}$ and $\phi_{\rm R}$ could in principle be a steady state: We could simply set $\chi_\alpha = \phi_\alpha$ for all sectors, and the $\phi_\alpha$'s would not change with time. This would, however, make it impossible to adjust the proteome's composition between day and night, as we have argued in the main text must occur, and it would moreover represent a fairly inefficient use of resources, with a substantial fraction of the C and R proteins typically inactivated. The qORAC model focuses on this second, efficiency concern and postulates that the $\chi_\alpha$'s should be chosen so that the system is always trying to move towards the highest growth rate that can be attained in its current, instantaneous environment.

Specifically, for any given environment, characterized by a ratio $\nu_{\rm C}/\nu_{\rm R}$, there exists an optimal partitioning of the proteome, $\phi_{\rm C}^{\rm opt}(\nu_{\rm C}/\nu_{\rm R})$, $\phi_{\rm R}^{\rm opt}(\nu_{\rm C}/\nu_{\rm R})$, and a corresponding optimal amino-acid level, $a^{\rm opt}(\nu_{\rm C}/\nu_{\rm R})$, that, together, maximize the instantaneous growth rate. This growth rate $\lambda^{\max}$ is obtained by maximizing the growth rate subject to the constraint

$$\phi_{\rm R} + \phi_{\rm C} = \phi_{\rm R,max}, \tag{S13}$$

with $\phi_{\rm R,max} = 1 - \phi_{\rm Q}$ the maximal ribosome fraction. Together with Eqs. S11 and S12, Eq. S13 gives a total of 3 equations equations with four unknowns, $\phi_{\rm R}, \phi_{\rm C}, \lambda, a$. When we add the fourth condition that the growth rate is maximal, all four variables are uniquely determined.

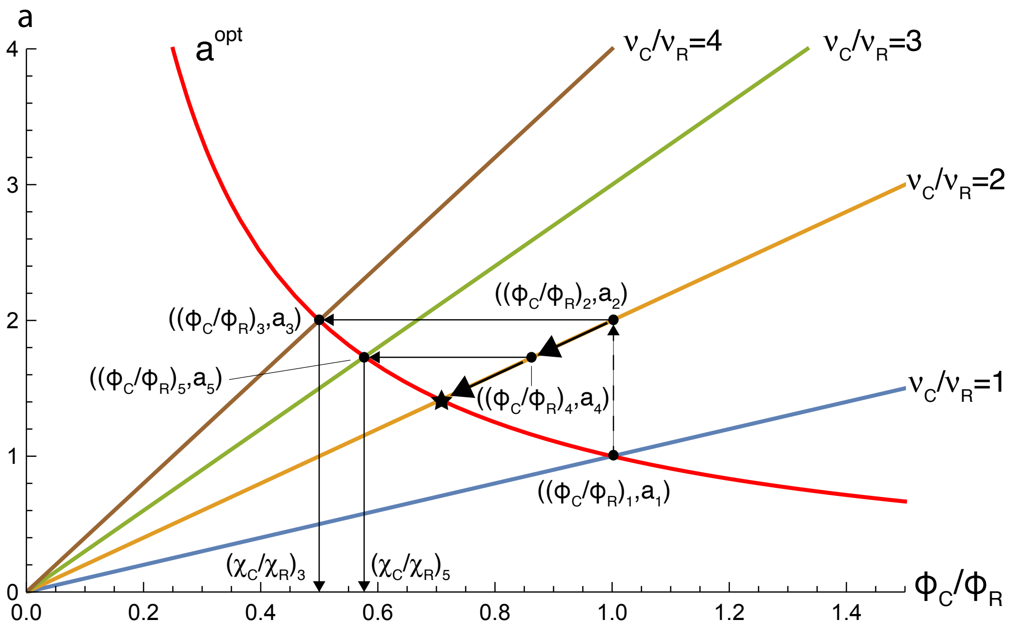


FIG. S1: **Illustration of the qORAC principle.** The qORAC principle illustrated for a an idealized, sudden change in the environment—a so-called "nutrient upshift" [1–3]—in which the system switches from a constant environment with $\nu_C/\nu_R = 1$ to another constant environment with $\nu_C/\nu_R = 2$. The blue, orange, green, and brown lines show the steady-state amino-acid concentration $a$ as a function of $\phi_C/\phi_R$ for four different environments, with $\nu_C/\nu_R = 1$ (environment $a$), $\nu_C/\nu_R = 2$ (environment $b$), $\nu_C/\nu_R = 3$ (environment $c$), and $\nu_C/\nu_R = 4$ (environment $d$), respectively; these lines represent solutions of Eq. S10 (which are straight lines for the specific parameters used in this figure but more generally might have some curvature). The intersections between the red curve and these steady-state lines give the system's optimal configuration $(\phi_C^{opt}/\phi_R^{opt}, a^{opt})$ that maximizes the growth rate for each environment $\nu_C/\nu_R$; the red curve is traced out parametrically as a function of $\nu_C/\nu_R$ by setting $\phi_C^{opt}/\phi_R^{opt}$ and $a^{opt}$ according to Eqs. S15-S17 for each value of $\nu_C/\nu_R$. Initially, the system is in the optimal configuration $((\phi_C/\phi_R)_1, a_1) = ((\phi_C^{opt}/\phi_R^{opt})_a, a_a^{opt})$ that maximizes the growth rate in environment $a$ with $\nu_C/\nu_R = 1$. Upon the shift to environment $b$ with $\nu_C/\nu_R = 2$, the amino-acid concentration instantly adjusts, and the configuration of the system becomes $((\phi_C/\phi_R)_2, a_2)$. The new concentration $a_2$ is not optimal for environment $b$ but is the optimal concentration $a_d^{opt}$ in another environment $d$, with $\nu_C/\nu_R = 4$, with an associated optimal protein partitioning $(\phi_C^{opt}/\phi_R^{opt})_d$; this is point $((\phi_C/\phi_R)_3, a_3) = ((\phi_C^{opt}/\phi_R^{opt})_d, a_d^{opt})$. According to qORAC, the system at point 2 allocates its ribosomes as if it were in this environment $d$; that is, it chooses the ribosome allocation $(\chi_C/\chi_R)_3 = (\phi_C^{opt}/\phi_R^{opt})_d$. Importantly, $(\chi_C/\chi_R)_3 = (\phi_C^{opt}/\phi_R^{opt})_d$ is lower than $(\phi_C/\phi_R)_2 = (\phi_C^{opt}/\phi_R^{opt})_a$, so according to Eq. S28 $\phi_C/\phi_R$ decreases: the system moves along the orange line towards the new fixed point, labeled with a star. At some later time, it will have arrived at point $((\phi_C/\phi_R)_4, a_4)$ with amino-acid concentration $a_4$. This concentration is equal to the optimal one $a_c^{opt}$ in environment $c$ with $\nu_C/\nu_R = 3$, with associated optimal protein partitioning $(\phi_C^{opt}/\phi_R^{opt})_c$, marked by the point $((\phi_C/\phi_R)_5, a_5) = ((\phi_C^{opt}/\phi_R^{opt})_c, a_c^{opt})$. The system again allocates its ribosomes to different protein sectors as if it were in this optimal configuration, leading to $(\chi_C/\chi_R)_5 = (\phi_C^{opt}/\phi_R^{opt})_c$, and as $(\chi_C/\chi_R)_5 < (\phi_C/\phi_R)_4$, the system continues to relax towards the starred fixed point. This point is the optimal configuration $((\phi_C^{opt}/\phi_R^{opt})_b, a^{opt})_b)$ that maximizes the growth rate in the new environment $b$. Parameters: $K_C = K_R = 1$; $\phi_{R,max} = 1$.

We then find

$$\lambda^{max} = \hat{\nu}_R \phi_R^{opt} = \hat{\nu}_C \phi_C^{opt}, \tag{S14}$$

$$\phi_R^{opt} = \frac{\hat{\nu}_C}{\hat{\nu}_C + \hat{\nu}_R} \phi_{R,max}, \tag{S15}$$

$$\phi_C^{opt} = \frac{\hat{\nu}_R}{\hat{\nu}_C + \hat{\nu}_R} \phi_{R,max}, \tag{S16}$$

$$a^{opt} = \sqrt{\frac{\nu_C}{\nu_R} K_R K_C}, \tag{S17}$$

where, for brevity of notation, we have introduced

$$\hat{\nu}_R \equiv \nu_R \alpha^\beta \frac{a^{opt}}{K_R + a^{opt}}, \tag{S18}$$

$$\hat{\nu}_C \equiv \nu_C \alpha^\beta \frac{K_C}{K_C + a^{opt}}. \tag{S19}$$

As these optimal quantities are defined instantaneously at each time point, they are not intrinsically time-dependent, but if $\nu_C = \nu_C(t)$ depends on time, as it may in our circadian system, then all of the optimal quantities inherit time dependence from $\nu_C(t)$.

The question then becomes: how does the system steer itself towards the optimal configuration $(\phi_R^{opt}(t), \phi_C^{opt}(t), a^{opt}(t))$? To answer this question, we note that if the nutrient conditions do not change with time, *i.e.* $\nu_{\rm C}(t) = \nu_{\rm C}$, and the system is in steady state in the optimal configuration that maximizes the growth rate, such that Eqs. S14-S17 hold, then Eq. S17 provides a unique mapping between the amino-acid concentration and the nutrient condition $\nu_{\rm C}/\nu_{\rm R}$. In other words, under the assumption that the growth rate is maximized, the steady state amino acid level $a$ provides an internal representation of the external environment described by the $\nu_\alpha$'s. The key assumption of qORAC is that the system uses the current concentration $a(t)$ to infer a value for $\nu_{\rm C}(t)/\nu_{\rm R}$ via the mapping Eq. S17, even when the system is not (yet) in its optimal steady state, and the inferred $\nu_{\rm C}(t)/\nu_{\rm R}$ thus does not equal the actual, current value of this ratio. The system then it allocates its ribosomes as if it were (already) growing in steady state in the optimal configuration with the inferred $\nu_{\rm C}(t)/\nu_{\rm R}$. That is, the system uses the steady state rule to translate from $a(t)$ to its desired ribosome allocations, whether or not it is actually in steady state; somewhat magically, doing this turns out to always drive the system towards the optimal steady state.

More specifically, in steady state it is possible to write the $\phi^{\rm opt}$'s as a function only of $a^{\rm opt}$, so that the system may determine the $\phi^{\rm opt}$'s simply by measuring its free amino acid levels, without ever explicitly calculating the ratio $\nu_{\rm C}/\nu_{\rm R}$. We first rewrite Eq. S16 so that it clearly depends only on the ratio of the $\nu$'s,

$$\phi_{\rm C}^{\rm opt} = \frac{1}{\hat{\nu}_{\rm C}/\hat{\nu}_{\rm R}+1}\phi_{\rm R,max} \tag{S20}$$

$$= \frac{\frac{a^{\rm opt}}{K_{\rm R}+a^{\rm opt}}}{\left(\frac{\nu_{\rm C}}{\nu_{\rm R}}\right)\frac{K_{\rm C}}{K_{\rm C}+a^{\rm opt}}+\frac{a^{\rm opt}}{K_{\rm R}+a^{\rm opt}}}\phi_{\rm R,max}\ , \tag{S21}$$

and similarly for $\phi_{\rm R}^{\rm opt}$. Inverting Eq. S17, we also have

$$\frac{\nu_{\rm C}}{\nu_{\rm R}} = \frac{(a^{\rm opt})^2}{K_{\rm R}K_{\rm C}}\ . \tag{S22}$$

Substituting this expression into Eq. S21 yields

$$\frac{\phi_{\rm C}^{\rm opt}(a^{\rm opt})}{\phi_{\rm R,max}} = \frac{\frac{a^{\rm opt}}{K_{\rm R}+a^{\rm opt}}}{\left(\frac{(a^{\rm opt})^2}{K_{\rm R}K_{\rm C}}\right)\frac{K_{\rm C}}{K_{\rm C}+a^{\rm opt}}+\frac{a^{\rm opt}}{K_{\rm R}+a^{\rm opt}}}\ , \tag{S23}$$

where all explicit dependence on the $\nu$'s has been eliminated. A similar procedure can be applied to $\phi_{\rm R}^{\rm opt}$. Eq. S23 thus shows that Eq. S16 maybe be rewritten so that the $\phi_{\rm C}^{\rm opt}$ is a function only of $a^{\rm opt}$, without any explicit dependence on the $\nu$'s. This rewriting of Eq. S16 into Eq. S23 enables the key step of qORAC that gives the ribosome allocation fractions $\chi_\alpha$ from which the temporal evolution of the protein sectors then follows, namely to replace $a^{\rm opt}$ in this function by the instantaneous amino acid level $a(t)$:

$$\chi_{\rm R}(t) = \phi_{\rm R}^{\rm opt}[a(t)], \tag{S24}$$

$$\chi_{\rm C}(t) = \phi_{\rm C}^{\rm opt}[a(t)]. \tag{S25}$$

These equations sum up the working principle of qORAC: the system interprets the current amino-acid level $a(t)$ as if it were already in the steady state that maximizes the growth rate, thus assuming $a(t) = a^{\rm opt}$; the system then allocates its ribosomes based on the sizes $\phi_\alpha^{\rm opt}$ of the protein sectors in that presumed optimal, steady state, $\chi_\alpha(t) = \phi_\alpha^{\rm opt}$.

We note that $a(t)$ is itself determined at each instant by the ratio $\phi_{\rm C}(t)/\phi_{\rm R}(t)$ through Eq. S10. Thus, we may rewrite Eqs. S24 and S25 as

$$\chi_{\rm R}(t) = \phi_{\rm R}^{\rm opt}\left[a\left(\frac{\phi_{\rm C}(t)}{\phi_{\rm R}(t)}\right)\right] \tag{S26}$$

$$\chi_{\rm C}(t) = \phi_{\rm R}^{\rm opt}\left[a\left(\frac{\phi_{\rm C}(t)}{\phi_{\rm R}(t)}\right)\right]\ , \tag{S27}$$

where the function $a(\phi_{\rm C}(t)/\phi_{\rm R}(t))$ is determined by solving Eq. S10 for $a(t)$. Eqs. S26 and S27, together with Eqs. 21 - 22 of the main text, constitute a closed pair of evolution equations for $\phi_{\rm R}(t)$ and $\phi_{\rm C}(t)$.

Perhaps surprisingly, in a constant environment characterized by a given value of $\nu_{\rm C}/\nu_{\rm R}$, these equations always steer the system towards the optimal configuration $(\phi_{\rm R}^{\rm opt}(t), \phi_{\rm C}^{\rm opt}(t), a^{\rm opt}(t))$ that maximizes its growth rate. That this must be the case is illustrated in Fig. S1, which follows the time evolution of the ratio $\phi_{\rm C}(t)/\phi_{\rm R}(t)$: Because $\phi_{\rm R}(t)+\phi_{\rm C}(t) = \phi_{\rm R,max}$ at all times, our dynamical system in fact only has a single independent, dynamical variable, which we can take to be $\phi_{\rm C}(t)/\phi_{\rm R}(t)$. Using Eqs. 21-22 from the main text for the time evolution of $\phi_{\rm R}$ and $\phi_{\rm C}$, we immediately find that this variable obeys

$$\frac{d[\phi_{\rm C}(t)/\phi_{\rm R}(t)]}{dt} = (\lambda\chi_{\rm R}/\phi_{\rm R})[(\chi_{\rm C}/\chi_{\rm R}) - (\phi_{\rm C}/\phi_{\rm R})], \tag{S28}$$

implying that the dynamics always drives $\phi_{\rm C}(t)/\phi_{\rm R}(t)$ towards $\chi_{\rm C}(t)/\chi_{\rm R}(t)$. The figure then shows that, for constant $\nu_{\rm C}/\nu_{\rm R}$, there is only a single fixed point where $\chi_{\rm C}/\chi_{\rm R} = \phi_{\rm C}/\phi_{\rm R}$, which is at the optimum, and qORAC's choice of $\chi_{\rm C}(t)/\chi_{\rm R}(t)$ always moves $\phi_{\rm C}(t)/\phi_{\rm R}(t)$ towards this fixed point. Importantly, the qualitative shape of the curves in the figure, and thus these conclusions, depends only on two facts, both of which hold for all parameter values. First, $a(t)$ is an increasing function of $\nu_{\rm C}(t)\phi_{\rm C}(t)/\nu_{\rm R}\phi_{\rm R}(t)$ (Eq. S10); and, second, $a^{\rm opt}$ is an increasing function of $\nu_{\rm C}/\nu_{\rm R}$ (Eq. S17).

Thus, given $\nu_{\rm C}(t)$, $\nu_{\rm R}$, $\phi_{\rm R}(t)$, and $\phi_{\rm C}(t)$, the qORAC model provides a prescription that uniquely determines the growth rate $\lambda(t)$ and protein synthesis fractions $\chi_{\rm R}(t)$ and $\chi_{\rm C}(t)$, and thereby the time evolution of the system. This time evolution tries to maximize the system's growth rate in any given environment. For general parameter choices, however, the values of $\lambda, \chi_{\rm R}, \text{and} \chi_{\rm C}$ specified by qORAC are quite complicated functions of the $\nu$'s and $\phi$'s. We show in the next section how this complexity reduces to a simple, physically satsifying picture in an appropriate limit.

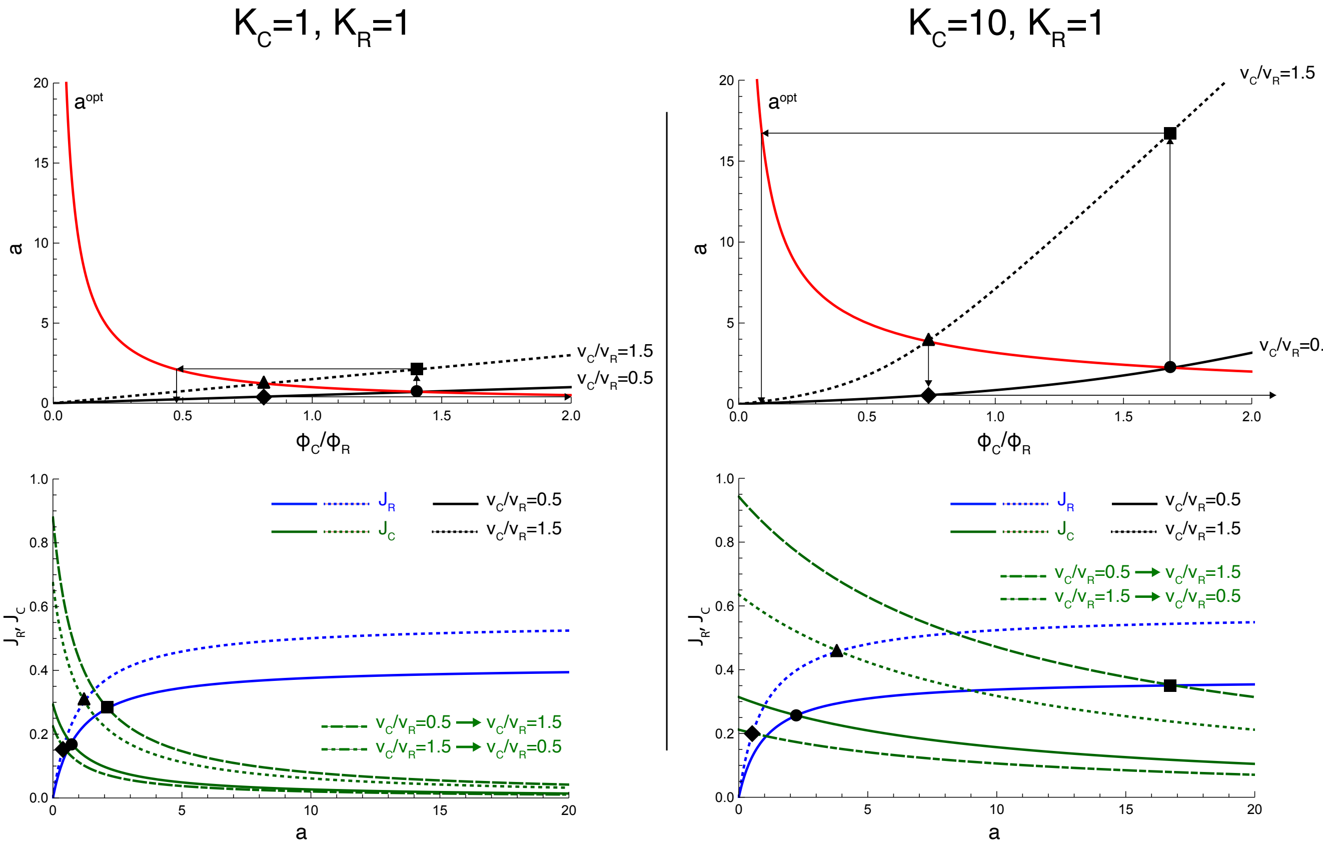


FIG. S2: **Importance of $K_{\rm C}/K_{\rm R}$ for the response to a nutrient upshift and downshift.** The panels in the left column corresponds to $K_{\rm C} = K_{\rm R} = 1$, while those in the right column correspond to $K_{\rm C} = 10, K_{\rm R} = 1$. The top panels show the same type of curves as Fig. S1. The bottom panels show the fluxes $J_{\rm C}$ (green) and $J_{\rm R}$ (blue) through the carbon and ribosome sectors, respectively, as a function of the amino-acid concentration $a$. The solid lines and dotted lines correspond respectively to $\nu_{\rm C}/\nu_{\rm R} = 0.5$ and to $\nu_{\rm C}/\nu_{\rm R} = 1.5$ in the steady state, after the protein mass fractions $\phi_{\rm C}$ and $\phi_{\rm R}$ have relaxed to their optimal values. In contrast, the long-dashed green line showes $J_{\rm C}$ immediately after an instantaneous nutrient upshift from $\nu_{\rm C}/\nu_{\rm R} = 0.5$ to $\nu_{\rm C}/\nu_{\rm R} = 1.5$, when the sectors have not had the time to adjust, while the dot-dashed green line shows $J_{\rm C}$ for the reverse transition. (We assume that the shifts occur through sudden changes in $\nu_{\rm C}$, leaving $\nu_{\rm R}$ fixed, so that $J_{\rm R}$ is unchanged immediately after a shift.) The filled circles, squares, triangles, and diamonds, in both the top and bottom panels, correspond to the quasi-equilibirum condition where the fluxes through the ribosome and carbon sectors are balanced and Eq. S10 is obeyed. The flux $J_{\rm R}$ through the ribosomal sector is given by Eq. S6 and the flux $J_{\rm C}$ through the carbon sector is given by Eq. S5 with, for the solid and dotted lines, $(\phi_{\rm C}, \phi_{\rm R})$ given by the optimal $(\phi_{\rm C}^{\rm opt}, \phi_{\rm R}^{\rm opt})$ for the respective environments, $\nu_{\rm C} = 0.5, \nu_{\rm R} = 1$ and $\nu_{\rm C} = 1.5, \nu_{\rm R} = 1$; $J_{\rm C}$ for the nutrient upshift (green dashed line) is computed with $\nu_{\rm C} = 1.5, \nu_{\rm R} = 1$ but $(\phi_{\rm C}, \phi_{\rm R})$ by the optimal values in the original environment $\nu_{\rm C} = 0.5, \nu_{\rm R} = 1$, and conversely for the downshift (green dot-dashed line); $\alpha^{\beta} = 1$. The top panels show that increasing $K_{\rm C}/K_{\rm R}$ generates a stronger response in the amino-acid concentration and protein allocation fraction $\chi_{\rm C}/\chi_{\rm R}$ for both a nutrient upshift and a downshift. The bottom panels illustrate that increasing $K_{\rm C}/K_{\rm R}$ ensures that during a nutrient shift the weakest link is always fully active, while the balance of fluxes is maintained by downregulating the activity of the strongest link. More specifically, a larger $K_{\rm C}$, which weakens the repression of the carbon sector, generates during a nutrient upshift a stronger increase in the amino-acid concentration and hence a stronger activation of the ribosome sector, which is the weakest link upon a nutrient upshift; in the limit that $K_{\rm C} \gg K_{\rm R}$ the ribosome sector is indeed fully active upon the nutrient upshift. Upon a downshift, the stronger drop in the amino-acid concentration helps to lift the repression of the carbon sector, which is now the weakest link. Note also that upon a nutrient downshift, the fluxes and hence the growth rate always first decrease, but then (partially) recover because the protein sectors adjust, as observed for *E. coli* [2]. Comparing the bottom panels also shows that increasing $K_{\rm C}$ increases the steady-state growth rate in both environments, because of the weaker repression of the carbon sector; in fact, it can be shown mathematically that in the limit that $K_{\rm C} \gg K_{\rm R}$ both sectors are fully active in steady state under constant nutrient conditions, thus fully maximizing the growth rate.

##### B. A simple limiting case

The reader may have noticed that, within the qORAC model just described, the active fractions of the carbon and ribosome sectors are always less than their maximal values: $\alpha_{\rm R}(t) < \alpha^{\beta}$ and $\alpha_{\rm C}(t) < \alpha^{\beta}$, even when the system has reached steady state in constant conditions. This would seem to imply that the qORAC model cannot relax at long times to the quasi-equilibrium model described in the main text, where $\alpha_{\rm R}(t) = \alpha_{\rm C}(t) = \alpha^{\beta}$. It turns out, however, that we can circumvent this difficulty when $K_{\rm R} \ll K_{\rm C}$. Then there is a large range of values of $a$ for which $K_{\rm R} \ll a \ll KC$, and both of the Michaelis-Menten functions in Eqs. S7 and S8 are close to unity. In particular, in steady state, according to Eq. S17, $a^{\rm opt}$ is of order $\sqrt{K_R K_C}$, and $K_R \ll a^{\rm opt} \ll K_C$. To leading order in small $K_{\rm R}/K_{\rm C}$, Eqs. S7 and S8 then reduce to $\alpha_{\rm R} = \alpha_{\rm C} = \alpha^{\beta}$, and Eq. S9 reduces to $\nu_{\rm C}\phi_{\rm C} = \nu_{\rm R}\phi_{\rm R}$. Thus, for any $\nu_{\rm C}/\nu_{\rm R}$, the steady state of the qORAC model agrees with the quasi-equilibrium model to a good approximation when $K_{\rm R} \ll K_{\rm C}$.

The limit $K_{\rm R} \ll K_{\rm C}$ also leads to simple behavior away from steady state. To see this analytically, we start by rewriting Eq. S9 in terms of the rescaled variables $\epsilon \equiv K_{\rm R}/K_{\rm C}$, $x \equiv a/K_{\rm C}$, and $r \equiv \nu_{\rm C}\phi_{\rm C}/(\nu_{\rm R}\phi_{\rm R})$ as

$$\frac{r}{1+x} = \frac{x}{\epsilon + x}\ . \tag{S29}$$

The arguments of the preceding paragraph imply that the system eventually relaxes to a steady state where $r = 1$ (up to small corrections that vanish as $\epsilon \to 0$.) Away from the steady state, however, $r$ will not be approximately 1. It is then easy to see that, in the limit of small $\epsilon$, the (unique) positive value of $x$ that solves Eq. S29 has different behavior for $r > 1$ and $r < 1$. Indeed, the expression $\frac{r}{1+x}$ on the lefthand side decreases from a maximum value of $r$ until it approaches zero for values of $x$ of order unity. On the other hand, the expression $\frac{x}{\epsilon+x}$ on the righthand side increases from zero to an asymptotic value of 1 that it approaches for $x$ values of order $\epsilon$. Thus, the two curves intersect at an $x$ value of order 1 when $r > 1$ and at an $x$ value of order $\epsilon$ when $r < 1$. (Since Eq. S29 is a quadratic equation in $x$, this behavior can of course also be checked directly from the exact solution.)

Translating these conclusions back into the original variables, we have found that when $\nu_{\rm C}\phi_{\rm C} > \nu_{\rm R}\phi_{\rm R}$ (and continuing to assume $K_{\rm R} \ll K_{\rm C}$ or $\epsilon \ll 1$), we need $a \sim K_{\rm C}$ in order to balance the fluxes in and out of the amino acid pool. Then, according to Eqs. S7 and S8, $\alpha_{\rm R} \approx 1$ while $\alpha_{\rm C} < 1$. That is, we downregulate the flux through the carbon sector so that it does not exceed the maximum flux possible through the ribosome sector. Conversely, when $\nu_{\rm C}\phi_{\rm C} < \nu_{\rm R}\phi_{\rm R}$, we need $a \sim K_{\rm R}$ to balance the fluxes, and then $\alpha_{\rm C} \approx 1$ while $\alpha_{\rm R} < 1$: We downregulate the flux through the ribosome sector so that it does not exceed the maximum flux possible through the carbon sector.

The upshot is that, to leading order in the limit $K_{\rm R} \ll K_{\rm C}$, the growth rate at all times $t$ is determined by the smaller of the two maximum fluxes:

$$\lambda(t) = \min\{\lambda_{\rm R}(t), \lambda_{\rm C}(t)\}, \tag{S30}$$

where $\lambda_{\rm R}$ and $\lambda_{\rm C}$ are the maximum fluxes through the ribosome and carbon sectors, respectively, and correspond to Eqs. 2-3 in the main text in the special case that $\phi_{\rm SC} = \phi_{\rm O} = 0$:

$$\lambda_{\rm R}(t) \equiv \alpha^{\beta}\nu_{\rm R}\phi_{\rm R}(t), \tag{S31}$$

$$\lambda_{\rm C}(t) \equiv \alpha^{\beta}\nu_{\rm C}(t)\phi_{\rm C}(t). \tag{S32}$$

The expressions for the ribosome allocation fractions $\chi_{\alpha}$ also simplify in the limit $K_{\rm R} \ll K_{\rm C}$. If $\nu_{\rm C}\phi_{\rm C} > \nu_{\rm R}\phi_{\rm R}$, then $a$ is of order $K_{\rm C}$. Following the prescription of Eq. S25 to obtain $\chi_{\rm C}$, we can substitute an $a$ value of order $K_{\rm C}$ into Eq. S23, discovering that $\chi_{\rm C}/\phi_{\rm R,max}$ is then of order $\epsilon = K_{\rm R}/K_{\rm C}$. The analogous calculation for $\chi_{\rm R}$ shows that $\chi_{\rm R}/\phi_{\rm R,max} = 1$, up to corrections of order $\epsilon$. Similarly, when $\nu_{\rm C}\phi_{\rm C} < \nu_{\rm R}\phi_{\rm R}$, it is a straightforward exercise to show that $\chi_{\rm C}/\phi_{\rm R,max} = 1$, up to corrections of order $\epsilon$, while $\chi_{\rm R}/\phi_{\rm R,max}$ is of order $\epsilon$. Hence, to leading order in small $\epsilon$, all available ribosomes are allocated to whichever sector has the smaller maximum flux:

$$\text{If } \lambda_{\rm C}(t) < \lambda_{\rm R}(t):\ \chi_{\rm C}(t) = \phi_{\rm R,max} \text{ and } \chi_{\rm R}(t) = 0 \tag{S33}$$

$$\text{If } \lambda_{\rm C}(t) > \lambda_{\rm R}(t):\ \chi_{\rm C}(t) = 0 \text{ and } \chi_{\rm R}(t) = \phi_{\rm R,max} \tag{S34}$$

In words, when the carbon sector limits growth, i.e. $\lambda_{\rm C}(t) < \lambda_{\rm R}(t)$, all ribosomes are allocated to upregulating the carbon sector, while if the ribosome sector is limiting, all ribosomes are devoted to upregulating the ribosome sector.

The effect of taking $K_{\rm C} \gg K_{\rm R}$ can also be understood graphically for the simple case of a step change in the environment (Fig. S2). Adapting terminology usually applied to non-photosynthetic bacteria [1–3], we refer to a sudden increase in $\nu_{\rm C}$ as a "nutrient upshift" and a sudden decrease in $\nu_{\rm C}$ as a "nutrient downshift". The repression threshold of the carbon sector $K_{\rm C}$ cannot be infinite in order to prevent the uncontrolled accumulation of amino acids during a strong nutrient upshift in which $\nu_{\rm C} > \nu_{\rm R}$. Yet, as the bottom panels highlight, to maximize the strength of the response, $K_{\rm C}$ should be larger than the activation threshold $K_{\rm R}$ of the ribosome sector such that during a nutrient upshift the amino-acid concentration rises sufficiently to fully activate the ribosome sector, the weakest link. Conversely, during a nutrient downshift, the limit $K_{\rm C} \gg K_{\rm R}$ ensures that the carbon sector, which is now the weakest link, is fully active. By fully activating the weakest protein sector, the growth rate is thus maximized on short timescales, shortly after the nutrient change. Yet, as the top panels show, the strong response of the amino-acid concentration also leads to a strong response of the ribosome allocation ratio $\chi_{\rm C}/\chi_{\rm R}$, which speeds up the adaptation of the protein sectors. And finally, as can be deduced from the bottom panels and shown mathematically, the limit $K_{\mathrm{C}} \gg K_{\mathrm{R}}$ also causes both sectors to be fully active under constant nutrient conditions, thus maximizing the growth rate in the long-time limit. We thus argue that the parameter regime $K_{\mathrm{C}} \gg K_{\mathrm{R}}$ is advantageous both for responding to nutrient upshifts and downshifts and for steady state growth.

In summary, then, in the limit $K_{\mathrm{R}} \ll K_{\mathrm{C}}$, the qORAC model follows two simple rules: First, the growth rate $\lambda(t)$ is set by the growth rate associated with the limiting sector, i.e. by the smaller of $\lambda_{\mathrm{R}}(t)$ or $\lambda_{\mathrm{C}}(t)$. Second, newly synthesized amino acids are, to the maximum extent possible, allocated to the limiting sector: If $\lambda_{\mathrm{R}} < \lambda_{\mathrm{C}}$, then $\chi_{\mathrm{C}} = 0$, so that all available amino acids go to the R sector, and the limiting growth rate can increase as quickly as possible (Eq. S34); similarly, if $\lambda_{\mathrm{C}} < \lambda_{\mathrm{R}}$, then $\chi_{\mathrm{R}} = 0$, and all available amino acids go to the C sector (Eq. S33). These rules generalize straightforwardly to the model considered in the main text that also includes a carbon storing sector $\phi_{\mathrm{SC}}$ and an OPP sector $\phi_{\mathrm{O}}$, giving rise to Eqs. 23-28, thereby illustrating a possible mechanistic basis for the phenomenological arguments used to justify these equations in the main text.

#### II. DEPENDENCIES ON $\phi_{\mathrm{O}}$

##### A. $\phi^*_{\mathrm{O}}$ in the anticipation model

In the main text, we systematically optimized over the target size $\phi^*_{\mathrm{SC}}$ of the carbon storing fraction and over the night-time active fraction $\alpha^{\mathrm{D}}$ to find the best possible growth rate for each value of the anticipation time $T_a$. One variable that we did not consider during this optimization process is the target size $\phi^*_{\mathrm{O}}$ of the OPP (Oxidative Pentose Phosphate) sector at night. In our model, this sector represents the set of proteins that must be expressed during the night to maintain cellular function in the absence of light. Although, in principle, $\phi^*_{\mathrm{O}}$ is a free parameter that we should optimize over to obtain the best possible performance, we show here that doing so yields at best only a very modest improvement in the overall growth rate. This justifies our choice to omit optimization over this parameter in our main results.

This point is illustrated in Fig. S3 (compare Fig. 4 of the main text), where we study the behavior of the anticipation model for different values of $\phi^*_{\mathrm{O}}$. As the figure demonstrates, the overall growth rate exhibits only minor changes in response to variations in $\phi^*_{\mathrm{O}}$, particularly near the optimal anticipation time, and especially when compared to the much larger effects seen when optimizing over parameters such as $\phi^*_{\mathrm{SC}}$, $T_a$, or $\alpha^{\mathrm{D}}$ (compare, e.g., Figs. 2–4 in the main text).

The selected values of $\phi^*_{\mathrm{O}}$ determine the corresponding optimal values of $\phi^*_{\mathrm{SC}}$, $\alpha^{\mathrm{D}}$, and $T_a$. The anticipation time $T_a$ is of particular interest: In the anticipation model at the optimal parameter values, the actual mass fraction $\phi_{\mathrm{O}}$ does not usually approach $\phi^*_{\mathrm{O}}$ at night. (See, e.g., Fig. 4E in the main text, where the blue curve shows that the maximal nighttime value of $\phi_{\mathrm{O}}$ is around 0.07, even though $\phi^*_{\mathrm{O}} = 0.15$.) Instead, $\phi^*_{\mathrm{O}}$ controls how fast $\phi_{\mathrm{O}}$ increases at the end of the day (through Eq. 22 of the main text, with $\chi_O = \phi^*_{\mathrm{O}}$ once the anticipation time has been reached), and a faster increase can be compensated by allowing a shorter time $T_a$ for the increase to occur, as seen in the figure. By trading off increases in $\phi^*_{\mathrm{O}}$ and decreases in $T_a$, the system can reach essentially the same nighttime level of $\phi_{\mathrm{O}}$, which is determined by the need to meet the $\eta$ constraint. The optimal growth rate then remains largely unchanged, confirming that optimization over $\phi^*_{\mathrm{O}}$ has a limited impact compared to the primary control variables.

##### B. Constant $\phi_{\mathrm{O}}$ model

In our discussion of the slow relaxation model in the main text, we noted that an alternative strategy, which we called the constant $\phi_{\mathrm{O}}$ model, is also open to a cell that is unable to anticipate. Here, we present a few results about this model.

We define the constant $\phi_{\mathrm{O}}$ model to be identical to the slow relaxation model, except that, whereas in the slow relaxation model the target OPPP sector size $\phi^*_{\mathrm{O}}$ is zero during the day and takes a constant value at night, in the constant $\phi_{\mathrm{O}}$ model, $\phi^*_{\mathrm{O}}$ takes the same constant value during the day and during the night, so that in steady state the actual OPPP mass fraction $\phi_{\mathrm{O}}$ is also constant at all times and is equal to $\phi^*_{\mathrm{O}}$. As we noted in the main text, the strategy embodied by this model has the advantage that the cell has enough OPPP sector proteins to carry out its nighttime tasks even while it does not grow at night, but the disadvantage that a fraction of ribosomes are devoted to holding $\phi_{\mathrm{O}}$ steady early in the day when they could more productively be employed to make proteins that contribute to exponential growth.

Fig. S4A shows the maximum average growth rate in the constant $\phi_{\mathrm{O}}$ model as a function of $\phi_{\mathrm{O}}$. This growth rate is found by optimizing over $\phi^*_{\mathrm{SC}}$ for each value of $\phi_{\mathrm{O}}$, subject to the constraint that the correct fluct $\eta$ passes through the OPPP at night; we set $\alpha^{\mathrm{D}}$ identically equal to zero, since the whole point of the constant $\phi_{\mathrm{O}}$ model is to circumvent the need to grow at night. One can see from Fig. S4A that the average growth rate depends more strongly on $\phi_{\mathrm{O}}$ in the constant $\phi_{\mathrm{O}}$ model than it does in the anticipation model (Fig. S3). The highest growth rate is attained when $\phi_{\mathrm{O}} = 0.07$ is close to the smallest possible value. For smaller values of $\phi_{\mathrm{O}}$, it quickly becomes necessary to produce large amounts of excess glycogen (frefconstphiofigC and F) so as to saturate the Hill function in Eq. 9 of the main text throughout the night and thus be able to meet the $\eta$ constraint of Eq. 8. By the time $\phi_{\mathrm{O}}$ has decreased to $\phi_{\mathrm{O}} = 0.05$, it is impossible to meet this constraint even when $g(t) \to \infty$. For values of $\phi_{\mathrm{O}}$ above the optimum at $\phi_{\mathrm{O}} = 0.07$, the

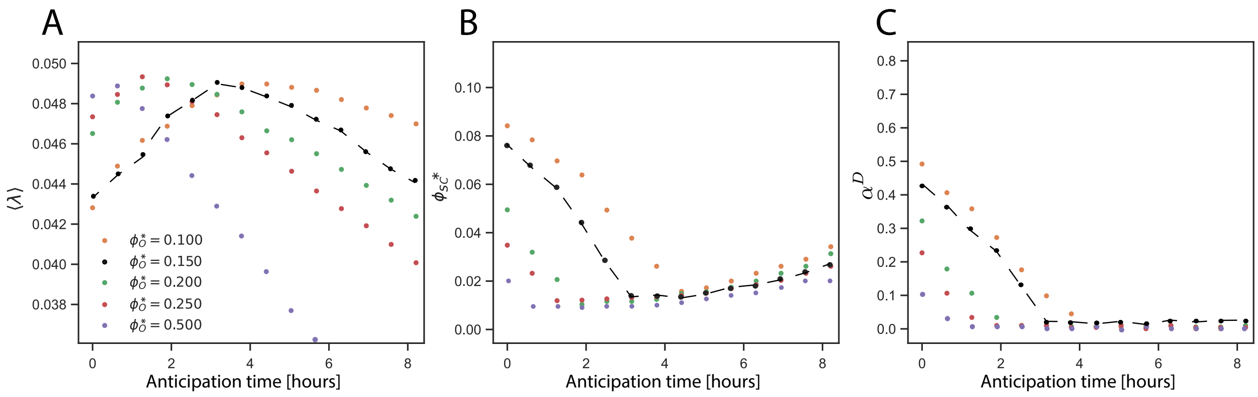


FIG. S3: Anticipation Model optimization for different values of $\phi_{\mathrm{O}}^*$ under the constraint of $\eta = 0.0975$. Panel A shows that the maximal average growth rate $\langle\lambda\rangle$, obtained by optimizing over the anticipation time and $\phi_{\mathrm{SC}}^*$ and $\alpha^{\mathrm{D}}$, only weakly depends on $\phi_{\mathrm{O}}^*$. The black dots, highlighted with a dashed line in panel A, corresponds to the value of $\phi_{\mathrm{O}}^*$ used in the main text. Panels B and C show, respectively, the optimal values of $\phi_{\mathrm{SC}}^*$ and $\alpha^{\mathrm{D}}$ that maximize the average growth rate over 24 hrs as a function of the anticipation time. Note the limited range of values of $\langle\lambda\rangle$ on the vertical axis in panel A.

growth rate decreases essentially linearly with $\phi_{\mathrm{O}}$, with a slope that is very close to the decrease in growth rate per unit increase in the expression level $\phi_{\mathrm{Q}}$ of inert protein (see Fig. S7 below). In other words, the excess $\phi_{\mathrm{O}}$ acts as a drag on growth equivalent to that of expressing inert protein. In the anticipation model, changes in $\phi_{\mathrm{O}}^*$ can be compensated by changes in the anticipation time $T_a$, so that, for a range of different values of $\phi_{\mathrm{O}}^*$, there is just enough OPPP protein present at nightfall to meet the $\eta$ constraint. Thus, the growth rate does not depend strongly on $\phi_{\mathrm{O}}^*$. In contrast, in the constant $\phi_{\mathrm{O}}$ model, any increase in $\phi_{\mathrm{O}}$ corresponds directly to an increase in the total amount of OPPP protein that the cell must produce, and any protein beyond the minimum amount needed to meet the $\eta$ constraint is essentially wasted, causing an appreciable decrease in the overall growth rate.

Quantitatively, it can be seen from Figs. S3 and S4 that the constant $\phi_{\mathrm{O}}$ model performs somewhat better than the slow relaxation model, but that both are outperformed by the anticipation model. The average growth rates are $\langle\lambda\rangle = 0.044\mathrm{h}^{-1}$ for the slow relaxation model; $\langle\lambda\rangle = 0.0471\mathrm{h}^{-1}$ for the constant $\phi_{\mathrm{O}}$ model at the optimal value of $\phi_{\mathrm{O}} = 0.07$; $\langle\lambda\rangle = 0.0484\mathrm{h}^{-1}$ for the anticipation model with $\phi_{\mathrm{O}}^* = 0.15$ as in the main text; and $\langle\lambda\rangle = 0.0493\mathrm{h}^{-1}$ for the anticipation model with $\phi_{\mathrm{O}}^* = 0.25$, where the growth rate turns out to be slightly higher. Hence, the average growth rate for the anticipation model is about 10% higher than for the slow relaxation model and about 5% higher than for the constant $\phi_{\mathrm{O}}$ model.

#### III. PRINCIPLE MASS HARMONICS ANALYSIS (PMHA) METHOD

In this section, we provide some more details on the steps used to arrive at the PMHA results reported in Fig. 5 of the main text.

Ref. [4] provides the measured abundances of each protein $i$ in the form of a time series $\tilde{X}_i(t)$ of the log ratio of the instantaneous abundance at time $t$ to the average abundance. Because these log ratios are better behaved than the actual abundances, we choose to begin our analysis on them, expanding each one in a Fourier series that we truncate after the first harmonic:

$$\tilde{X}_i(t) \simeq \tilde{A}_i + \tilde{a}_i \sin(\omega t) + \tilde{b}_i \cos(\omega t), \tag{S35}$$

where we assume a common oscillation frequency $\omega = 2\pi/24\,\mathrm{h}$ for all proteins.

To convert the log ratios into actual expression ratios relative to the mean, we apply the transformation:

$$\hat{X}_i = e^{\tilde{X}_i} \simeq e^{\tilde{A}_i + \tilde{a}_i \sin(\omega t) + \tilde{b}_i \cos(\omega t)}. \tag{S36}$$

Here, the log ratio $\tilde{X}_i$ represents the natural logarithm of the ratio between the average copy number per cell and the instantaneous copy number at a given time. Since this representation inherently implies that $\tilde{A}_i$ is small, and given that log-ratio fluctuations are typically moderate, the coefficients $\tilde{a}_i$ and $\tilde{b}_i$ are also expected to be small perturbations. Under these conditions, we expand the exponential function to first order:

$$\hat{X}_i \simeq e^{\tilde{A}_i}[1 + \tilde{a}_i \sin(\omega t) + \tilde{b}_i \cos(\omega t)], \tag{S37}$$

Consequently, we obtain:

$$\hat{X}_i \simeq \hat{A}_i + \hat{a}_i \sin(\omega t) + \hat{b}_i \cos(\omega t), \tag{S38}$$

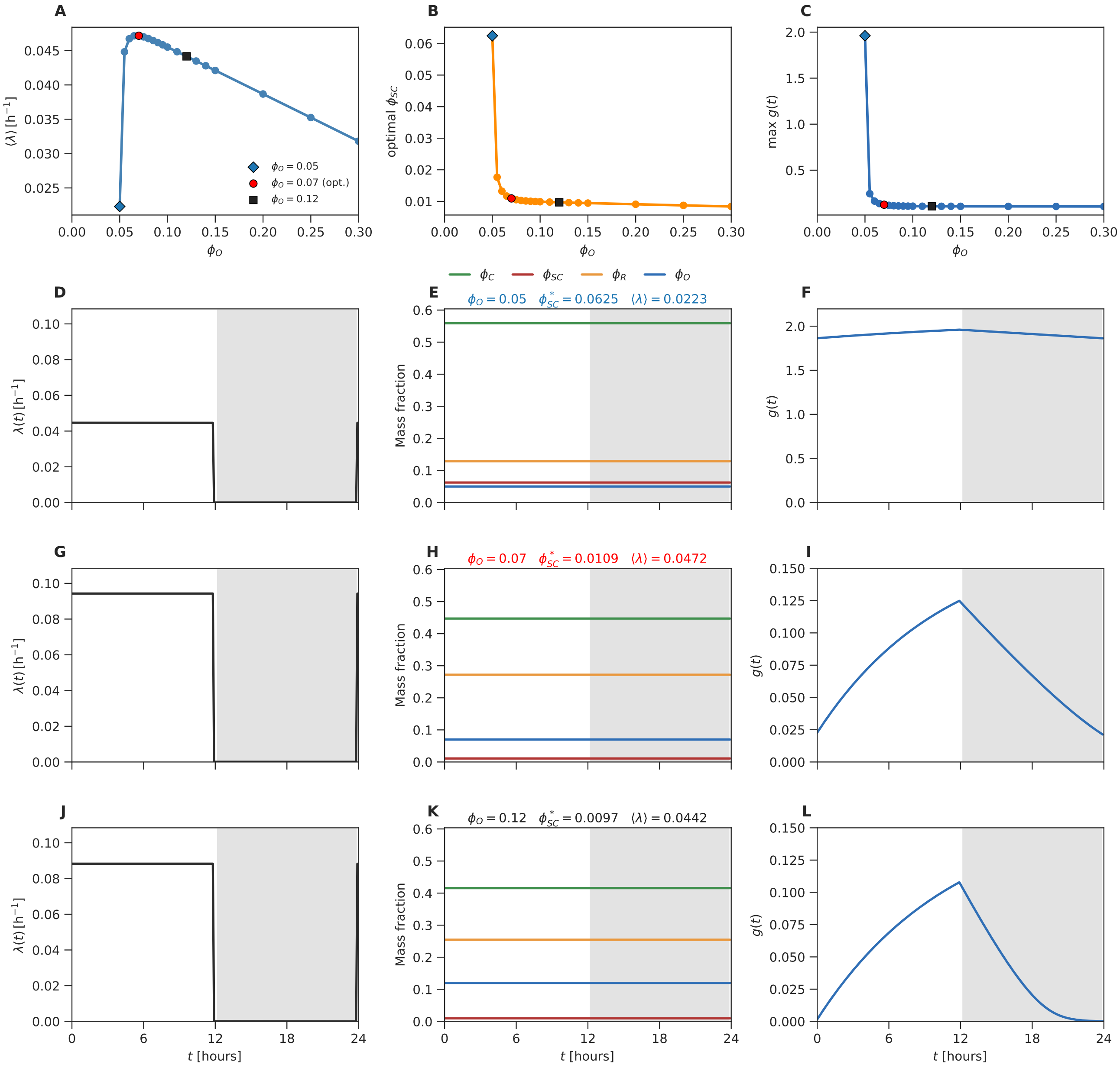


FIG. S4: The constant $\phi_{\rm O}$ model. (A) Average growth rate over 24h as a function of $\phi_{\rm O}$. The maximum growth rate occurs at $\phi_{\rm O} = 0.07$ (red dot). Below that value, the growth rate drops off rapidly as it becomes necessary to store large quantities of excess glycogen in order to keep the flux through the OPP pathway (Eq. 8 of the main text) at the correct value $\eta$. Above $\phi_{\rm O} = 0.07$, the growth rate decreases linearly with $\phi_{\rm O}$, consistent with the idea that increasing the $\phi_{\rm O}$ level above the minimum needed to consume almost all glycogen while meeting the $\eta$ constraint is essentially the same as expressing excess inert protein (Fig. S7 below). The blue diamond and black square mark representative points with $\phi_{\rm O}$ below and above the optimum value, respectively. (B) Optimal value of $\phi^*_{\rm SC}$, as a function of $\phi_{\rm O}$. (C) Maximum value of the stored glycogen level $g(t)$ over the 24h cycle, generally corresponding to the value at $t = 12$h, as a function of $\phi_{\rm O}$. (D-L) Time traces for the three representative points marked in panel A. (D-F), blue diamond, $\phi_{\rm O} - 0.05$; (G-I) red dot at the optimal value, $\phi_{\rm O} = 0.07$; (J-L) black square, $\phi_{\rm O} = 0.12$. The left, middle, and right columns give, respectively, the instantaneous growth rate $\lambda(t)$, the sector fractions $\phi_\alpha(t)$, and the stored glycogen level $g(t)$. Note that all of the ribosome allocations $\chi_\alpha$ (main text Eqs. 20-22) are constant in this model during the day, and there is no growth at night; as a consequence, all of the sector fractions $\phi_\alpha$ are strictly constant.

where the new coefficients are defined as $\hat{A}_i = e^{\tilde{A}_i}$, $\hat{a}_i = e^{\tilde{A}_i}\tilde{a}_i$, and $\hat{b}_i = e^{\tilde{A}_i}\tilde{b}_i$.

Ref. [4] also separately provides the time-averaged number of copies of each protein in an experimental sample of fixed overall protein mass. When multiplied by the protein molecular masses, these give a figure proportional to the time-averaged mass fraction of each protein. To obtain the final protein mass fractions $X_i(t)$ (Eq. 29 of the main text), we scale the ratios $\hat{X}_i$ so that $\hat{A}_i$ equals this time average.

#### IV. METHODS

##### A. Model parameters

In choosing model parameters, we sought to arrive at biologically reasonable numbers informed by available measurements on *S. elongatus*. (We note, however, that available data are not sufficient to support a well-conditioned fitting procedure that would reliably arrive at unique "best-fit" values for all of the parameters.) To this end, we first estimate values for the parameters associated with the largest fluxes in our model, through the C and R sectors, which most directly control the growth rate, while ignoring the effects of smaller perturbations associated with other sectors. Once the key parameters associated with the C and R sectors have been set, we then use additional observations to set the remaining parameters associated with glycogen storage and the OPPP sector.

Specifically, in our model, the key parameters that can be varied experimentally are $\nu_{\mathrm{C}}^{\mathrm{L}}$, which is determined by the $\mathrm{CO}_2$ and light levels during the day, and $\nu_{\mathrm{R}}$, which can be varied experimentally via a translational inhibitor such as chloramphenicol. We begin by setting the parameter $\nu_{\mathrm{R}}$, because that determines the maximum growth rate. This maximum rate is reached at saturating $\mathrm{CO}_2$ and light intensity levels, which in our model corresponds to the limit $\nu_{\mathrm{C}}^{\mathrm{L}} \to \infty$. The highest growth rate measured for *S. elongatus* is that for the strain UTEX 2973. It can achieve a maximum doubling time of 1.5 h, compared to 4.9 h for the clock model strain PCC 7942 [5]. By replacing three alleles of strain PCC 7942 by the fast-growth associated alleles identified for UTEX 2973, the doubling time of PCC 7942 could be reduced to 2.3 h [6]. We imagine that this is the highest growth rate that the model strain PCC 7942 can achieve, and we therefore set $\nu_{\mathrm{R}}$ such that with $\nu_{\mathrm{C}}^{\mathrm{L}} \to \infty$, the cell doubling time is about 2 h. We then chose $\nu_{\mathrm{C}}^{\mathrm{L}}$ such that in the quasi-equilibrium model the doubling time under constant light (LL) conditions is 6 h, corresponding to a growth rate during the day $\lambda^{\mathrm{L}} = \ln(2)/6\mathrm{h}^{-1}$; this is consistent with relatively rapid growth under high light intensity [7]. Then, if there is minimal nighttime growth in LD conditions the doubling time in LD is about 12 h. This value of $\lambda^{\mathrm{L}}$, together with $\nu_{\mathrm{C}}^{\mathrm{L}}$ and $\nu_{\mathrm{R}}$, then also fixes $\phi_{\mathrm{C}}$ and $\phi_{\mathrm{R}}$ during the day in the quasi-equilibrium model through flux balance. In equations: $\lambda^{\max} = \ln(2)/2\mathrm{h}^{-1} = \nu_{\mathrm{R}}\phi_{\mathrm{R,max}}$, which sets $\nu_{\mathrm{R}}$, and $\lambda^{\mathrm{L}} = \ln(2)/6\mathrm{h}^{-1} = \nu_{\mathrm{C}}^{\mathrm{L}}\phi_{\mathrm{C}} = \nu_{\mathrm{R}}\phi_{\mathrm{R}}$, which, with $\phi_{\mathrm{R}} + \phi_{\mathrm{C}} = \phi_{\mathrm{R,max}}$, sets $\nu_{\mathrm{C}}^{\mathrm{L}}$, $\phi_{\mathrm{C}}$, and $\phi_{\mathrm{R}}$ in the quasi-equilibrium model during the day. In making these estimates, we ignore the small carbon flux that is devoted to storing glycogen [8] and, in the absence of a measured value for *S. elongatus*, set $\phi_{\mathrm{R,max}} = 0.8$, consistent with figures from *E. coli* [9].

The key parameters related to the production and consumption of glycogen, $\nu_{\mathrm{SC}}, \phi_{\mathrm{SC}}^*, \eta, \tilde{\nu}_{\mathrm{C}}^{\mathrm{D}}, K_{\mathrm{C}}, \tilde{\nu}_{\mathrm{O}}, \phi_{\mathrm{O}}*, K_{\mathrm{O}}$ are next set as follows. We eventually would like to systematically vary $\phi_{\mathrm{SC}}^*$ to find the optimal value that maximizes the growth rate over 24 hrs. Yet, this value will depend on $\nu_{\mathrm{SC}}$. To set $\nu_{\mathrm{SC}}$, we therefore first choose an interim value of $\phi_{\mathrm{SC}}^*$, which we will use to estimate $\nu_{\mathrm{SC}}$ and which will set the scale for the final value of $\nu_{\mathrm{SC}}$ found through the optimization procedure. Since the carbon-storing sector is small compared to the other sectors, we set, somewhat arbitrarily, this interim value to $\phi_{\mathrm{SC}}^* = 0.01$. To then set $\nu_{\mathrm{SC}}$, we exploit the observation that the peak mass fraction of glycogen per dry weight of cell mass is about 10% in the related cyanobacterium *Synechocystis* [8]. If we imagine that the glycogen mass ratio $g$, governed by Eqs. 15-16 of the main text, starts at zero at the beginning of the day and accumulates with $\nu_{\mathrm{SC}}\phi_{\mathrm{SC}} \approx \nu_{\mathrm{SC}}\phi_{\mathrm{SC}}^* \approx$ constant, then the concentration $T/2 = 12\mathrm{h}$ later at the end of the day is $g(T/2) = (\nu_{\mathrm{SC}}\phi_{\mathrm{SC}}^*/\lambda^{\mathrm{L}})[1-\exp(-\lambda^{\mathrm{L}}T/2)] = 0.75\nu_{\mathrm{SC}}\phi_{\mathrm{SC}}^*/\lambda^{\mathrm{L}}$, where we have used $\lambda^{\mathrm{L}} = \ln(2)/6\mathrm{h}^{-1}$ in the last equality. Setting this concentration equal to 0.1, and recalling that $\lambda^{\mathrm{L}} = \nu_{\mathrm{C}}^{\mathrm{L}}\phi_{\mathrm{C}}$, we conclude that $\nu_{\mathrm{SC}}\phi_{\mathrm{SC}}^* \approx 0.13\nu_{\mathrm{C}}^{\mathrm{L}}\phi_{\mathrm{C}}$. With $\phi_{\mathrm{C}}$ obtained from the quasi-equilibrium model above and $\phi_{\mathrm{SC}}^* = 0.01$, this equation specifies $\nu_{\mathrm{SC}}$.

To estimate $\eta$, we note that the integral of the third term in Eq. 16 of the main text from $T/2 = 12\mathrm{h}$ to $T = 24\mathrm{h}$ equals the flux constraint $\eta$. Moreover, if we assume that we are operating at (near)-optimal parameter values, then the cells do not grow at night ($\lambda(t) = 0$), and the second and fourth terms vanish; under the same assumptions, the glycogen concentration should be nearly zero by the end of the night, and the integral of this equation from $T/2 = 12\mathrm{h}$ to $T = 24\mathrm{h}$ yields $\eta = g(T/2) = g_{\max}$. Finally, we can then conclude from our solution of the same equation during the day in the preceding paragraph that $\eta = g_{\max} = (\nu_{\mathrm{SC}}\phi_{\mathrm{SC}}^*/\lambda^{\mathrm{L}})[1 - \exp(-\lambda^{\mathrm{L}}T/2)] = 0.75\nu_{\mathrm{SC}}\phi_{\mathrm{SC}}^*/\lambda^{\mathrm{L}}$.

The target size of the OPPP sector, $\phi_{\mathrm{O}}^*$, is set to 0.15, but we verify in Section II of this *Supporting Information* that our major conclusions are insensitive to the precise choice for $\phi_{\mathrm{O}}^*$. The maximal OPPP efficiency $\tilde{\nu}_{\mathrm{O}}$ is then chosen such that $\nu_{\mathrm{O}}\phi_{\mathrm{O}}^* 12\mathrm{h} > \eta \approx g_{\max}$. This choice—together with our choice of $K_{\mathrm{O}}$ discussed in the next paragraph—ensures that almost all glycogen is consumed before the end of the night. In practice, we have chosen $\tilde{\nu}_{\mathrm{O}}\phi_{\mathrm{O}}^* 12\mathrm{h} = 3\eta$. The precise value of $\tilde{\nu}_{\mathrm{O}}$ should not be important, because in the limit that all glycogen is consumed by the end of the night (which is the optimal strategy because then no glycogen goes to waste by dilution), the value of $\tilde{\nu}_{\rm O}$ only affects the moment during the night at which all glycogen is consumed, not how much must be stored during the day (which is controlled by $\eta$). To prevent big changes in the $\phi_{\rm C}$ and $\phi_{\rm R}$ sectors between day and night, the parameter for the carbon-processing efficiency at night is set to be equal to that during the day: $\tilde{\nu}_{\rm C}^{\rm D} = \nu_{\rm C}^{\rm L}$. This choice also ensures that our main results reflect generic advantages of the anticipation enabled by the presence of a clock and are not biased one way or the other by differences between $\tilde{\nu}_{\rm C}^{\rm D}$ and $\nu_{\rm C}^{\rm L}$.

The constants $K_{\rm C}$ and $K_{\rm O}$ are set such that at the beginning of the night, the OPPP sector $\phi_{\rm O}$ is saturated, meaning that $K_{\rm C}, K_{\rm O} < g_{\rm max}$; specifically, we take $K_{\rm C} = 0.1 g_{\rm max}$ and $K_{\rm O} = 0.2 g_{\rm max}$. As for $\tilde{\nu}_{\rm O}$, the precise values of $K_{\rm C}, K_{\rm O}$ should only affect how fast glycogen is consumed during the night, not how much needs to be stored during the day. In particular, we expect that, for optimal parameter values, $\alpha^{\rm D} \approx 0$; in this case, the value of $K_{\rm C}$ will be irrevelant at the optimum.

The final parameters values obtained through this procedure are listed in Table. S1. (As discussed above and in the main text, we optimize over the parameters $\phi_{\rm SC}^*$ and $\alpha^{\rm D}$ and thus do not assign them specific values here.)

##### B. Numerical methods

The optimization over $\alpha^{\rm D}$, $\phi_{\rm SC}^*$, and $T_a$ is performed by numerically propagating our model using a standard ode solver, and computing the average growth rate $\langle \lambda \rangle_T = 1/T \int_0^T \lambda(t)$ over one circadian period of duration $T =$ 24h. We take particular care to ensure that the model has been run for enough oscillation cycles that it has settled down to its asymptotic limit cycle before performing this average. In the numerical implementation of qORAC, we use the bang-bang response of Eqs. S33 and S34 when $\lambda_{\rm C}$ and $\lambda_{\rm R}$ differ by more than a tolerance factor of $5 \times 10^{-4}{\rm h}^{-1}$ and treat $\lambda_{\rm C}$ and $\lambda_{\rm R}$ as effectively equal if they differ by less than this amount.

| Parameter | Value |
|---|---|
| $\nu_{\rm C}^{\rm L}$ | 0.25 $\mathsf{h}^{-1}$ |
| $\nu_{\rm R}$ | 0.35 $\mathsf{h}^{-1}$ |
| $K_{\rm C}$ | 0.0098 |
| $K_{\rm O}$ | 0.0195 |
| $\tilde{\nu}_{\rm C}^{\rm D}$ | 0.25 $\mathsf{h}^{-1}$ |
| $\nu_{\rm SC}$ | 1.50 $\mathsf{h}^{-1}$ |
| $\tilde{\nu}_{\rm O}$ | 0.16 $\mathsf{h}^{-1}$ |
| $\phi_{\rm R,max}$ | 0.8 |
| $\phi_{\rm O}^*$ | 0.15 |
| $\eta$ | 0.0975 |

TABLE S1: Parameter values used in the model.

#### V. SUPPORTING FIGURES

Here we present additional supporting figures that provide complementary perspectives on the model's behavior, helping the reader to better appreciate its dynamics from different angles.

For the Slow Proteome Model, in Figs. S5 and S6 we provide additional plots to clarify the model's behavior at the three points marked on the heat map in Figure 3A of the main text (corresponding to three different values of $\phi_{\rm SC}^*$ and $\alpha$). Specifically, Fig. S5 replots the behavior of a number of variables of interest from Fig. 3 so that the time traces corresponding to the three points appear on the same axes and can be more easily compared. Particulary noteworthy is that in the optimal configuration that maximizes the growth rate, the glycogen concentration becomes zero precisely at the end of the night; at the suboptimal points 1 and 3, the glycogen concentration either never becomes zero (point 3) or becomes zero before the night is over (point 1).

Similarly, while the heat map of Figure 3A of the main text shows general trends, it is difficult to discern the behavior of the growth rate $\lambda$, $\phi_{\rm SC}^*$, and $\alpha$, along a contour line of constant $\eta$. To clarify this, in Fig. S6, we plot the behavior of the growth rate $\lambda$, the target carbon storage fraction $\phi_{\rm SC}^*$, and the activity $\alpha$, along the constant $\eta = 0.0975$ contour line highlighted in Fig. 3A.

Finally, Fig. S7 plots the maximum attainable growth rate in the anticipation model as a function of the maximum ribosome mass fraction $\phi_{\rm R,max}$. Because $\phi_{\rm R,max} = 1 - \phi_{\rm Q}$, where $\phi_{\rm Q}$ is the mass fraction of inert protein, this graph may equivalently be viewed as showing how the growth rate decreases as the expression of inert protein increases. This is of interest for at least two reasons. First, any clock is built from proteins whose expression does not contribute directly to growth and thus increases $\phi_{\rm Q}$; Fig. S7 indicates to what extent this extra protein expression counteracts the increase in growth rate due to anticipation that the clock enables. Second, a common technique to probe the physiology of microbial growth is observe how much the growth rate decreases as the experimentalist increases ectopic expression of an inert protein [9]. Fig. S7 predicts the results of such an experiment. It is worth underlining that the growth rates in the figure are optimized over the parameters $\alpha^{\rm D}$, $\phi_{\rm SC}^*$, and $T_a$ and thus more properly might be thought of as giving an upper bound on growth rates that could be measured experimentally. Rather than optimizing over three parameters in this way, one might be tempted instead to assume that no other parameters change their values as $\phi_{\rm R,max}$ is varied and thus, in particular, that $\alpha^{\rm D}$, $\phi_{\rm SC}^*$, and $T_a$ retain their optimal values from Fig. 4 of the main text. The difficulty with this approach is that changing only one parameter, like $\phi_{\rm R,max}$, generally also changes the nighttime flux $\eta$ through the OPPP sector; thus, to meet the $\eta$ constraint the cell must change some other parameter when it is forced to decrease $\phi_{\rm R,max}$. In the absence of a clear reason to choose one particular parameter for this role, we find it more informative to optimize over all of the free parameters.


[1] R. Planqué, J. Hulshof, B. Teusink, J. C. Hendriks, and F. J. Bruggeman, PLoS Computational Biology **14**, e1006412 (2018), ISSN 1553-734X.

[2] D. W. Erickson, S. J. Schink, V. Patsalo, J. R. Williamson, U. Gerland, and T. Hwa, Nature **551**, 119 (2017).

[3] M. Mori, S. Schink, D. W. Erickson, U. Gerland, and T. Hwa, Nature Communications pp. 1–7 (2017).

[4] A. C. L. Guerreiro, M. Benevento, R. Lehmann, B. v. Breukelen, H. Post, P. Giansanti, A. F. M. Altelaar, I. M. Axmann, and A. J. R. Heck, Molecular & Cellular Proteomics **13**, 2042 (2014), URL `https://www.ncbi.nlm.nih.gov/pmc/articles/PMC4125736/`.

[5] J. Ungerer, P.-C. Lin, H.-Y. Chen, and H. B. Pakrasi, mBio **9**, e02327 (2018), ISSN 2161-2129.

[6] J. Ungerer, K. E. Wendt, J. I. Hendry, C. D. Maranas, and H. B. Pakrasi, Proceedings of the National Academy of Sciences **115**, E11761 (2018), ISSN 0027-8424.

[7] S. W. Teng, S. Mukherji, J. R. Moffitt, S. de Buyl, and E. K. O'Shea, Science **340**, 737 (2013).

[8] T. Zavřel, M. Faizi, C. Loureiro, G. Poschmann, K. Stühler, M. Sinetova, A. Zorina, R. Steuer, and J. Červený, eLife **8**, e42508 (2019).

[9] M. Scott, C. W. Gunderson, E. M. Mateescu, Z. Zhang, and T. Hwa, Science **330**, 1099 (2010).

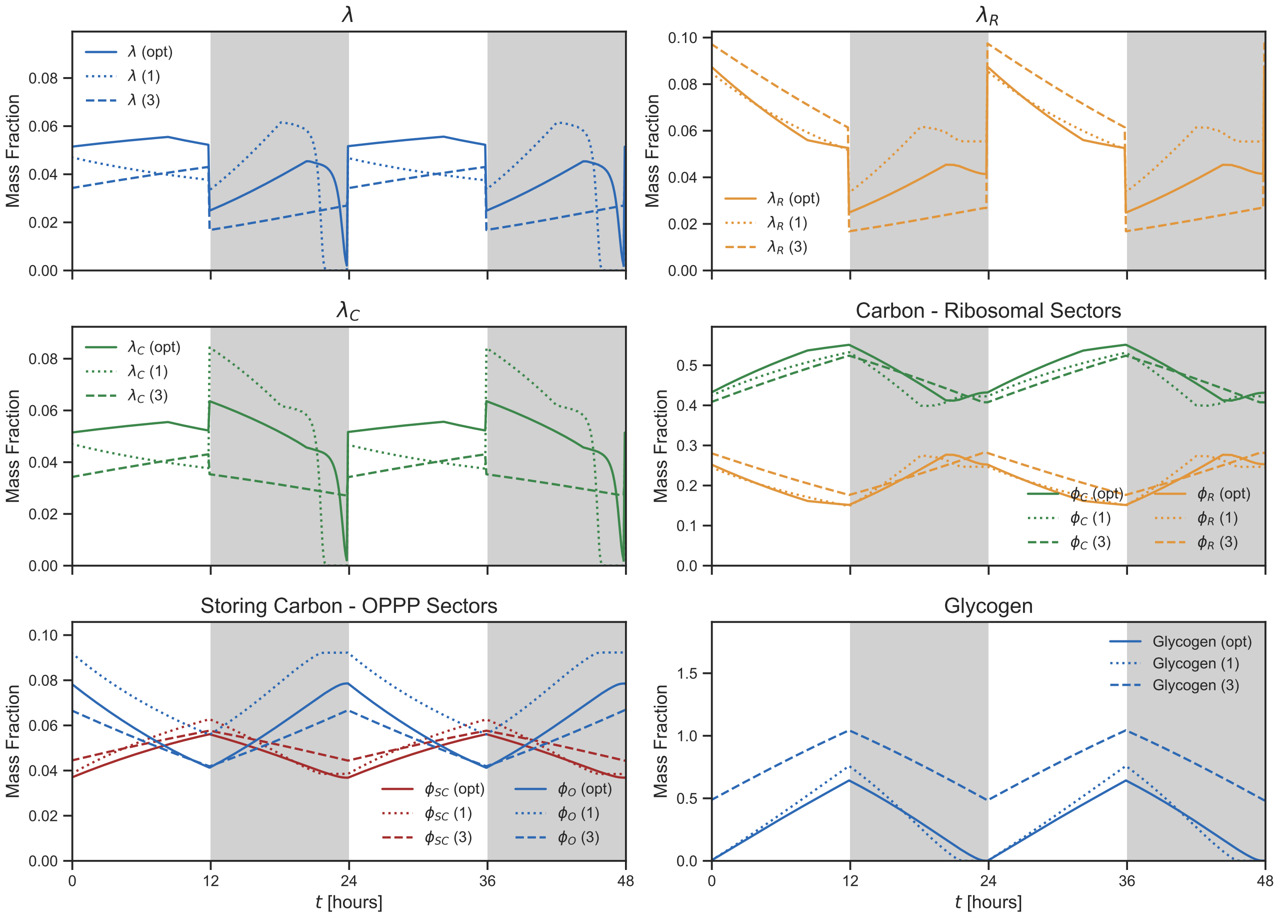


FIG. S5: Time traces of the Slow Proteome Model for different parameter configurations. The continuous line shows the behavior of the optimal model, whereas the dashed lines correspond to parameter sets chosen away from the optimum. In particular, we consider the parameter values associated with points 1 and 3 in Figure 3 of the main text. Plotting the mass fractions of the sectors and the time-dependent growth rate $\lambda(t)$ together highlights the differences in the system's behavior across configurations. Notably, the optimal model that maximizes the growth rate over 24 hrs stores and consumes glycogen in the most efficient way, avoiding both excessive accumulation and premature depletion, while suboptimal parameter choices lead to less balanced glycogen dynamics and reduced performance.

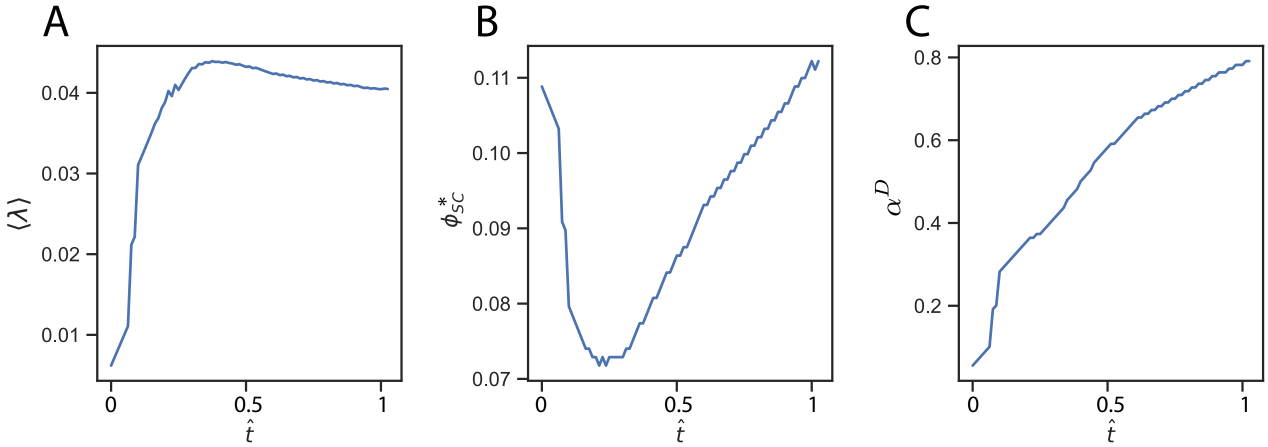


FIG. S6: Behavior of the Slow Proteome Model along the contour line of constant $\eta = 0.0975$ highlighted in Fig. 3A of the main text, with the progression along the contourline parametrized by the variable $\hat{t}$ that runs from 0 to 1 along the contour (with 0 corresponding to $\alpha^{\mathrm{D}} = 0$. Panel A: $\langle\lambda\rangle$. Panel B: $\phi^*_{\mathrm{SC}}$. Panel C: $\alpha^{\mathrm{D}}$. The maximal average growth rate along this contour line is $\langle\lambda\rangle = 0.044$, corresponding to $\phi^*_{\mathrm{SC}} = 0.077$ and $\alpha^{\mathrm{D}} = 0.473$.

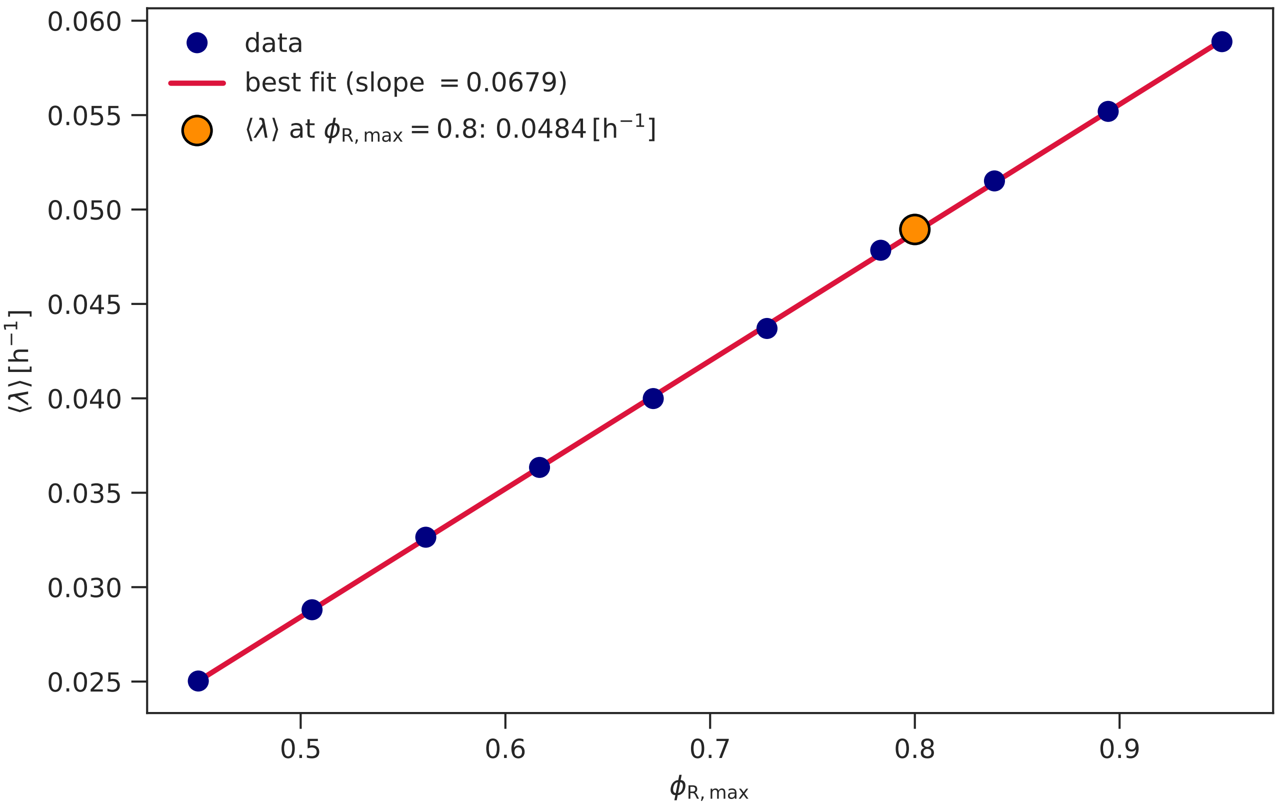


FIG. S7: Maximum achievable average growth rate $\langle\lambda\rangle$ as a function of the maximum possible mass fraction $\phi_{\mathrm{R,max}}$ of the ribosome sector, in the anticipation model. The orange dot indicates the growth rate at the value $\phi_{\mathrm{R,max}} = 0.8$ used throughout this paper, unless otherwise noted (see also Table. S1). For each value of $\phi_{\mathrm{R,max}}$ (black dots), we optimized the time-averaged growth rate over the parameters $\alpha^{\mathrm{D}}$ and $\phi^{*}_{\mathrm{SC}}$, subject to the constraint of fixed flux $\eta$ through the OPPP sector at night (Eq. 8 in the main text), and over the anticipation time $T_a$. Evidently, the dependence of $\langle\lambda\rangle$ on $\phi_{\mathrm{R,max}}$ is linear to a very good approximation, as also seen in steady state growth in *E. coli* [9]; the slope of the best-fit line (red) is $0.06785\mathrm{h}^{-1}$. Because the mass fraction of the inert sector is given by $\phi_{\mathrm{Q}} = 1 - \phi_{\mathrm{R,max}}$, decreasing $\phi_{\mathrm{R,max}}$ amounts to increasing the expression level $\phi_{\mathrm{Q}}$ of proteins that do not contribute to growth. This figure may thus be interpreted in two ways. First, it indicates the mass fraction of clock-associated proteins—which belong to $\phi_{\mathrm{Q}}$, because they are not directly involved in fixing carbon, protein synthesis, or similar core physiological functions—that can be tolerated before the extra protein expression overwhelms the increase in growth rate from clock-enabled anticipation. Comparing to Fig. 3 of the main text, we see that the maximum tolerable clock protein mass fraction for the growth rate in the anticipation model to remain above the growth rate in the slow proteome model is about 0.06, considerably above the mass fraction $\sim 0.002$ of the core Kai clock proteins in *S. elongatus* [4]. Second, the figure predicts how much the growth rate will decrease in experiments where extra inert proteins are expressed ectopically [9]. Importantly, in this latter interpretation the plot really gives an upper bound on what might be observed experimentally, because, faced with ectopic protein expression, the cell may not be able to optimize over all of the parameters that we have in our calculation.
